# In-vivo evidence for increased tau deposition in temporal lobe epilepsy

**DOI:** 10.1101/2025.05.14.654095

**Authors:** Raúl R. Cruces, Jack Lam, Thaera Arafat, Jessica Royer, Judy Chen, Ella Sahlas, Arielle Dascal, Daniel Mendelson, Raluca Pana, Robert Hopewell, Chris Hung-Hsin Hsiao, Gassan Massarweh, Jean-Paul Soucy, Pedro Rosa-Neto, Marie-Christine Guiot, Sylvia Villeneuve, Lorenzo Caciagli, Matthias J. Koepp, Andrea Bernasconi, Neda Bernasconi, Boris C. Bernhardt

**Affiliations:** McConnell Brain Imaging Centre, Montreal Neurological Institute, McGill University, Montreal, Québec, Canada; Montreal Neurological Institute, McGill University Health Centre (MUHC) and other McGill University; StoP-AD Centre, Douglas Mental Health University Institute, McGill University, Montréal, QC, Canada; UCL Queen Square Institute of Neurology, University College London, London WC1N 3BG, UK; Department of Neurology, Inselspital, Sleep-Wake-Epilepsy Center, Bern University Hospital, University of Bern, Bern, Switzerland

**Keywords:** Temporal lobe epilepsy, [^18^F]MK-6240, tauopathy, neuroimaging, neuroinflammation

## Abstract

Temporal lobe epilepsy (TLE), the most common pharmaco-resistant epilepsy in adults, has been linked to structural brain changes extending beyond the mesiotemporal areas. While not traditionally viewed as a neurodegenerative disorder, recent *ex-vivo* studies have shown elevated levels of misfolded tau protein in TLE. This study investigated tau deposition in TLE patients using the *in-vivo* PET tracer [18F]MK-6240. We studied 28 TLE patients and 28 healthy controls to assess tau uptake and its relationship with brain connectivity, clinical variables, and cognitive function alongside post-surgical tissue from a subset of patients. Compared to controls, TLE patients exhibited markedly increased [18F]MK-6240 uptake in bilateral superior and medial temporal regions and the parietal cortex, with tau accumulation following regional functional and structural connectivity and cognitive impairment. Immunohistochemistry analysis confirmed variable phosphorylated tau staining in 5/6 operated cases with available specimens. These findings suggest that tau accumulation contributes to cognitive decline observed in TLE, supporting a potential role of tau in epilepsy-related neurodegeneration.

## Introduction

Mounting evidence suggests that temporal lobe epilepsy (TLE), the most common pharmaco-resistant epilepsy in adults, is not a static condition, but one that is associated with progressive changes extending beyond the mesiotemporal disease epicenter^1–4^. Although the exact causes of disease progression remain unknown, several lines of evidence suggest similar mechanisms as in Alzheimer’s disease (AD), an established neurodegenerative condition. First, cognitive profiles in middle-aged pharmaco-resistant epilepsy patients resemble those of mild cognitive impairment, which can be linked to AD^5^. Furthermore, epidemiological studies have indicated an elevated risk for pharmaco-resistant epilepsy patients to be diagnosed with AD, and *vice versa*^6,7^. Across all age groups, patients with AD show a 3-fold higher risk of suffering from epilepsy^8,9^. Finally, a recent analysis of *ex-vivo* specimens in operated TLE patients has shown highly elevated levels of misfolded tau protein in operated TLE patients^10^, a hallmark of AD pathology^11^. Such findings complement data from animal models of TLE, which demonstrate that seizure activity triggers tau hyperphosphorylation and amyloidogenic pathways^12,13^.

Whole-brain patterns of increases of phosphorylated tau in epilepsy remain unknown due to limits imposed by the scope of resection. Also, while *ex-vivo* tau increases in TLE were related to postoperative cognitive decline^14^, correlations with measures of preoperative cognitive function remain incompletely understood. These considerations underscore the need for *in-vivo* analyses of tau in TLE to evaluate its potential role as both a modulator of cognitive function and a predictor of clinical outcomes. Research in AD and other neurodegenerative conditions has benefitted from the introduction of second-generation PET tracers that are sensitive and specific to pathological deposits. In particular, the [^18^F]-MK-6240 tracer (*henceforth* MK-6240) has shown high affinity and selectivity to intracellular neurofibrillary tangles^15–17^. In contrast, while there is an extensive PET literature in epilepsy on presurgical localization of hypometabolism via [^18^F]fluorodeoxyglucose^18,19^, PET studies in epilepsy have so far not systematically investigated tau tracers^7^. As such, the whole brain distribution of tau in epileptic patients remains unknown. Unlike research in AD, where tau uptake patterns are increasingly understood and contextualized with respect to patterns of brain connectivity^1–3,20–22^, there have not been any systematic assessments of potential network-level effects underlying the accumulation of these pathological deposits in TLE.

The objective of our study was to investigate the distribution and prevalence of tau aggregates in TLE using *in-vivo* MK-6240 PET. Tau mapping was complemented with network decoding models informed by multimodal MRI investigations in the same participants that tested for spatial associations of tau uptake in patients with patterns of global and local connectivity. Moreover, linear models examined associations with cognitive function assessed outside the scanner and clinical measures from the patients presurgical examination.

## Results

We recruited 28 TLE patients (14 Females, age=36±12) and 28 healthy volunteers (14 Females, age=33±7). with no significant differences in age and sex distribution. The mean onset of epilepsy in TLE patients was 21±14 years, the mean duration of epilepsy was 15.3±11.5 years, and 29% had hippocampal atrophy (volume or asymmetry less than 1.5 standard deviations from controls). Further clinical features are listed in **Table 1**. Analyses are referenced to seizure onset and reported separately for the ipsilateral and contralateral hemispheres.

**Table 1.**
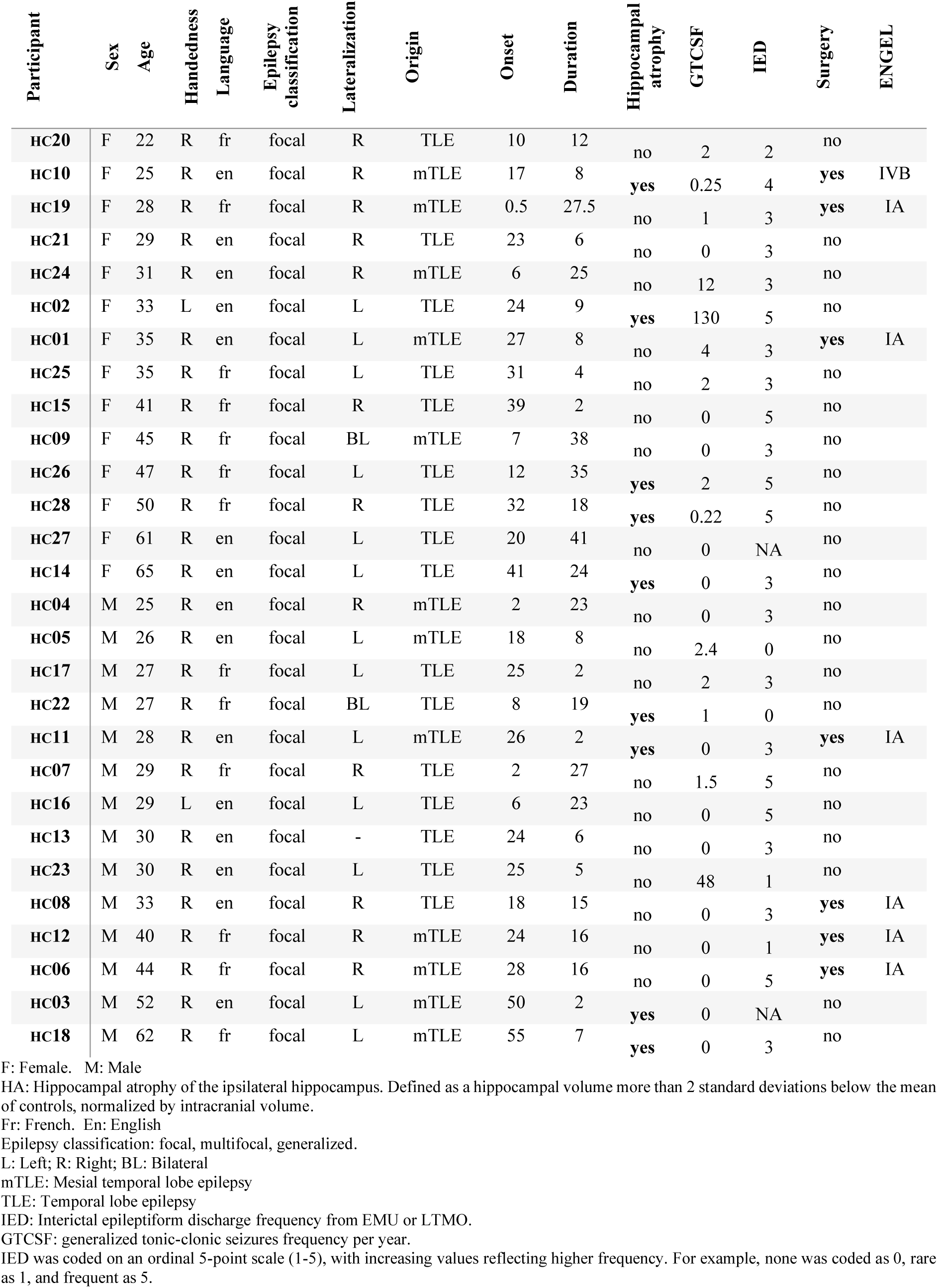
Clinical features.

### Widespread increases in tau uptake in TLE

Mean cortical MK-6240 SUVR was increased in patients with TLE relative to controls, both ipsilateral (t=3.16, p=<0.002) and contralateral (t=3.25, p=0.002) to the seizure focus. The individualized MK-6240 SUVR histogram analysis across the cortical surface of all subjects (**Figure 1A**) shows that there is a high prevalence of TLE patients with high MK-6240 uptake, which is reflected in the mean group uptake (**Figure 1B**). Indeed, 75% of TLE patients but no controls demonstrated elevated mean SUVR above 1.5 (97.5th percentile based on controls, **Table 2**), with a spatial distribution generally resembling the statistical effects. This pattern was more pronounced in the ipsilateral hemisphere but showed a similar bilateral distribution (**Supplementary Figure 1 and Table 2**).

**Figure 1.**
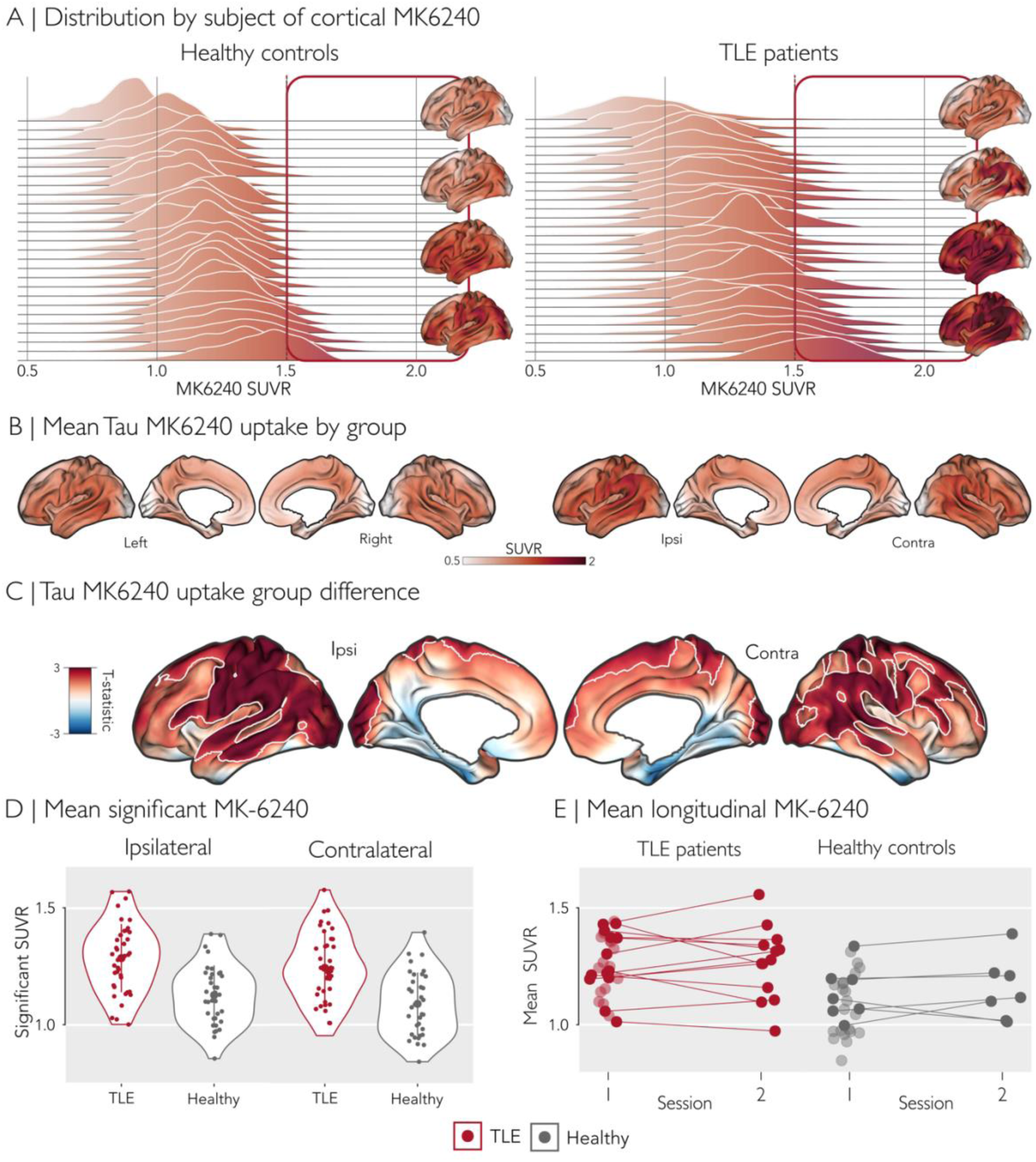
A) Subject-Wise distribution of cortical MK-6240 uptake. Density histograms illustrate the distribution of global cortical MK-6240 uptake for individual participants in each group. Red squares indicate regions where SUVR exceeds the abnormality threshold (SUVR > 97.5th percentile). Brain images represent individual participants from each group with progressively increasing MK-6240 values. **B) Group-wise mean tau MK-6240 uptake.** Cortical phosphorylated tau deposits were assessed with MK-6240 and visualized as standardized uptake value ratios (SUVR). The average MK-6240 uptake is shown for healthy controls and TLE patients. **C) Group differences in MK-6240 uptake.** Group comparisons between patients and controls were performed using a mixed-effects model, adjusting for age and sex, and corrected for multiple comparisons using random field theory (p_RFT_ < 0.025). Significant regions are outlined in white. **D) Mean SUVR values for significant regions** obtained from the group differences analysis. Violin plots are shown by hemisphere and group. **E) Longitudinal trajectories of MK-6240 uptake** by group between the PET scans 1 and 2. Lines correspond to the longitudinal participants, and transparent points are the cross-sectional subjects.

**Table 2.** SUVR thresholds.

| Group | 97.5 th | 99.0 th |
| --- | --- | --- |
| Healthy controls | 1.56 | 1.65 |
| All subjects | 1.69 | 1.82 |

Surface-wide analysis of the group difference of cortical MK-6240 SUVR indicated bilateral high uptake in patients relative to controls, particularly in superior-posterior temporal and parietal and occipital cortex (**Figure 1C**). Statistical comparisons were based on two-tailed mixed-effects models controlling for age and sex, with significance assessed using random field theory (p_RFT_<0.025). The strongest effects were localized to the ipsilateral frontal operculum and the posterior cortex, including the anterior and inferior parietal lobe (postcental, supramarginal and angular gyrus) and the lateral temporal (superior temporal gyrus) and occipital cortices. In the contralateral hemisphere, a similar pattern was observed, sparing the occipital region, but with a larger involvement of the temporal lobe, encompassing the posterior section of the superior and middle temporal gyri, and the premotor cortex.

In significant clusters derived from the main group-difference analysis, the overall effect was similar in both hemispheres but slightly higher ipsilateral to seizure onset (ipsilateral: t=5.04, p<0.001; contralateral: t=4.78, p<0.001, **Figure 1D**). No significant differences were observed in the subcortical structures analyzed; however we found a trend of higher MK-6240 SUVR uptake in the ipsilateral hippocampus (t=-2.00, uncorrected p =0.049, **Table 3**).

**Table 3.** Mean Subcortical MK-6240 SUVR values.

| *Structure* | *Patient* | *Healthy* | *T-statistic* | *p-value* |
| --- | --- | --- | --- | --- |
| Ipsilateral thalamus | 0.67±0.11 | 0.66±0.11 | 0.673 | 0.5 |
| Ipsilateral caudate | 0.63±0.12 | 0.62±0.12 | 0.845 | 0.4 |
| Ipsilateral putamen | 0.69±0.14 | 0.66±0.12 | 1.34 | 0.2 |
| Ipsilateral pallidus | 0.64±0.14 | 0.64±0.13 | 0.895 | 0.4 |
| Ipsilateral amygdala | 0.63±0.13 | 0.61±0.09 | 0.972 | 0.3 |
| Ipsilateral hippocampus | 0.71±0.12 | 0.71±0.11 | 0.412 | 0.7 |
| Ipsilateral accumbens | 0.59±0.14 | 0.60±0.17 | 0.411 | 0.7 |
| Contralateral thalamus | 0.69±0.10 | 0.65±0.10 | 1.27 | 0.2 |
| Contralateral caudate | 0.63±0.11 | 0.59±0.11 | 0.87 | 0.4 |
| Contralateral putamen | 0.68±0.12 | 0.63±0.13 | 1.5 | 0.14 |
| Contralateral pallidus | 0.62±0.14 | 0.57±0.13 | 0.718 | 0.5 |
| Contralateral amygdala | 0.64±0.10 | 0.60±0.11 | 1.28 | 0.2 |
| Contralateral hippocampus | 0.69±0.11 | 0.69±0.10 | -0.37 | 0.7 |
| Contralateral accumbens | 0.57±0.13 | 0.54±0.14 | 0.292 | 0.8 |

2 1 Mean±SD
3 2 Welch Two Sample t-test, **uncorrected** p-values for multiple comparisons.

To assess whether MK-6240 increases were seen even after correcting for grey matter atrophy, a common feature in TLE^23,24^, we re-ran the mixed effects model controlling vertex-wise for regional grey matter thickness estimates and our results were largely unaffected (**Supplementary Figure 2A-B**). Additionally, we evaluated the robustness of our results using different reference areas to calculate the MK-6240 SUVR and confirmed high consistency (**Supplementary Figure 3**).

We evaluated the effects of sex on MK-6240 uptake to investigate potential sex-related differences in tau pathology. Our findings indicated that females with TLE overall exhibited an elevated Tau burden relative to their male counterparts (**Supplementary Figure 4**).

A subgroup of 7 healthy controls and 13 TLE patients had longitudinal tau PET data available (mean difference between scan 1 and 2=18.75 months, minimum=12 months, maximum 42 months). While there was a non-significant trend for subtle increases in TLE in the ipsilateral temporal lobe at uncorrected thresholds (**Supplementary Figure 2C**), MK-6240 uptake showed no significant longitudinal change in either healthy controls or patients, and no group differences in rate of change were observed (**Figure 1E**).

### Association with demographic, clinical, and cognitive measures

Neither age MK-6240 SUVR (r = −0.03, p = 0.80) nor disease duration were significantly associated with MK-6240 SUVR in patients (r = −0.08, p = 0.68) in clusters of between-group differences. At the surface level, MK-6240 SUVR was associated with age in the ipsilateral medial-to-anterior cingulate gyrus and inferior frontal lobe, and with disease duration in the contralateral inferior parietal region and the antero-inferior portion of the occipital lobe. Notably, the latter duration association to tau remained consistent even after controlling for age. Age at seizure onset, contralateral hippocampal volume, interictal epileptiform discharge frequency and generalize tonic clonic seizure frequency were not correlated to MK-6240 SUVR, neither in clusters of between-group findings nor at the vertex-level (**Supplementary Figure 5**).

When studying associations to hippocampal volume, we noted that lower ipsilateral hippocampal volume related to higher MK-6240 SUVR in patients in clusters of findings (r = −0.28, p = 0.04; **Figure 2A-left**). Furthermore, lower ipsilateral hippocampal volumes were associated with higher MK-6240 SUVR in the contralateral posterior superior parietal lobe (**Figure 2A-right**).

**Figure 2.**
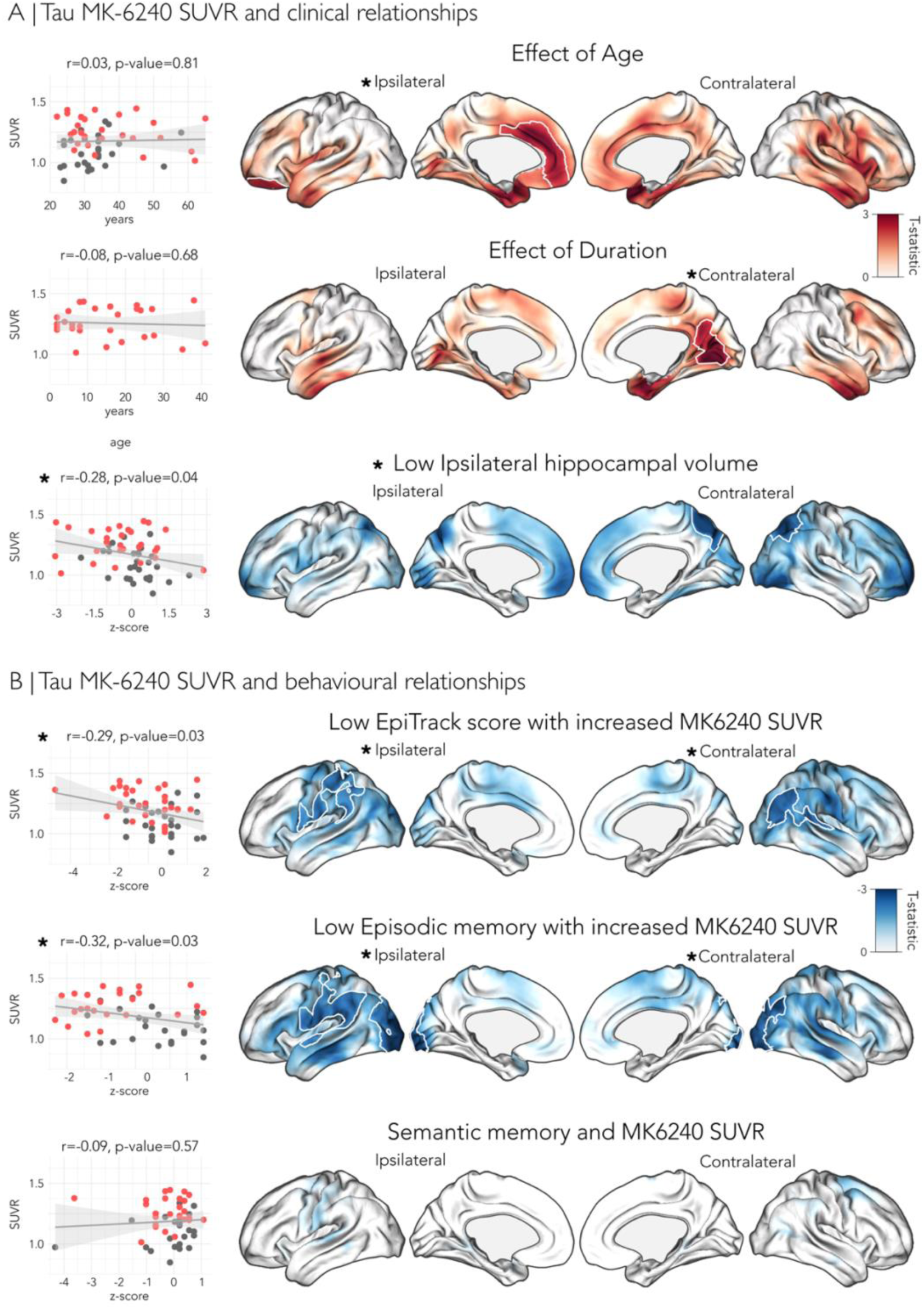
A) Tau MK-6240 SUVR and clinical relationships. Scatterplots show an association of mean MK-6240 SUVR in regions that showed significant between-group differences (see Figure 1) with clinical variables. Corresponding cortical projections visualize the regional effects of these variables on MK-6240 SUVR. Dark red on the brain maps indicate regions where age and duration are associated with increased tau deposition, as measured by t-statistics. **B) Tau MK-6240 SUVR and behavioral relationships.** Scatterplots display the relationship of mean MK-6240 SUVR in regions of significant-between group differences with behavioral measures. Brain maps illustrate the corresponding regional effects, dark blue indicating areas where lower cognitive performance is associated with higher MK-6240 SUVR. Mixed-effects models were applied to assess the association between MK-6240 SUVR and these features, corrected for multiple comparisons with random field theory (p<0.025, two tails). Significant regions after multiple comparisons are outlined in white in the surface-based model and an asterisk (*) indicates significant results in the analyses.

We also studied associations between clusters of tau PET measures and cognitive variables, notably scores on the EpiTrack^25^, as well as scores on previously validated tasks tapping into episodic and semantic memory^26^ (**Figure 2B-left**). In line with prior results^27^, our TLE patients showed lower performance on the *EpiTrack* (r = -0.29, p = 0.03) and *episodic memory task* (r = −0.32, p = 0.03), indicating an association between increased Tau burden and lower cognitive performance. In contrast, their performance did not differ from controls for the *semantic memory task* (*p*=0.09, p=0.57). At the surface level, cognitive performance measures had distinct associations with MK-6240 SUVR, lower EpiTrack scores showed only a trend toward higher MK-6240 SUVR, whereas reduced episodic memory performance exhibited a negative association with MK-6240 SUVR (**Figure 2B-right**). This pattern involved the ipsilateral angular gyrus and bilateral occipital lobes. No associations were observed for semantic memory (**Supplementary Figure 5**).

Additionally, we evaluate the impact of all available clinical features on MK-6240 uptake using a linear mixed-effects model. No significant clinical predictor evaluated in the model impacted MK-6240 uptake except for a larger patient–control difference in females than in males, and an association between a higher number of antiseizure medications and lower MK-6240 SUVR. We also assessed subcortical and hippocampal MK-6240 SUVR and did not observe any significant between group differences (**Supplementary Figure 6, Supplementary Table 3**). Moreover, we did not observe any significant correlations between subcortical MK-6240 SUVR and clinical or behavioral measures.

An exploratory subgroup analysis was performed comparing patients in the top 25 percent of SUVR uptake (n=8) with the remaining patients (n=20). While we did not find marked effect sizes for differences in hippocampal volumes, cognitive measures, seizure frequency, or interictal epileptiform discharges, we observed that age at onset was lower in the top 25 percent group compared to the remaining patients (13.38±10.20 vs 24.68±14.23, d>0.8). Moreover, the top 25 percent group showed lower path length derived from the structural connectome (0.13±0.09 vs 0.34±0.26; d>0.8), indicating altered network organization in patients with higher tau SUVR values.

### Association to network architecture

We assessed the susceptibility of globally highly interconnected regions to tauopathy using group-based network decoding models (**Figure 3**). Vertex-wise measures of mean connectivity and neighborhood-weighted connectivity informed by MK-6240 were derived from group-average structural and functional connectomes and correlated with vertex-wise *t*-statistics obtained from the group-difference mixed-effects model.

**Figure 3.**
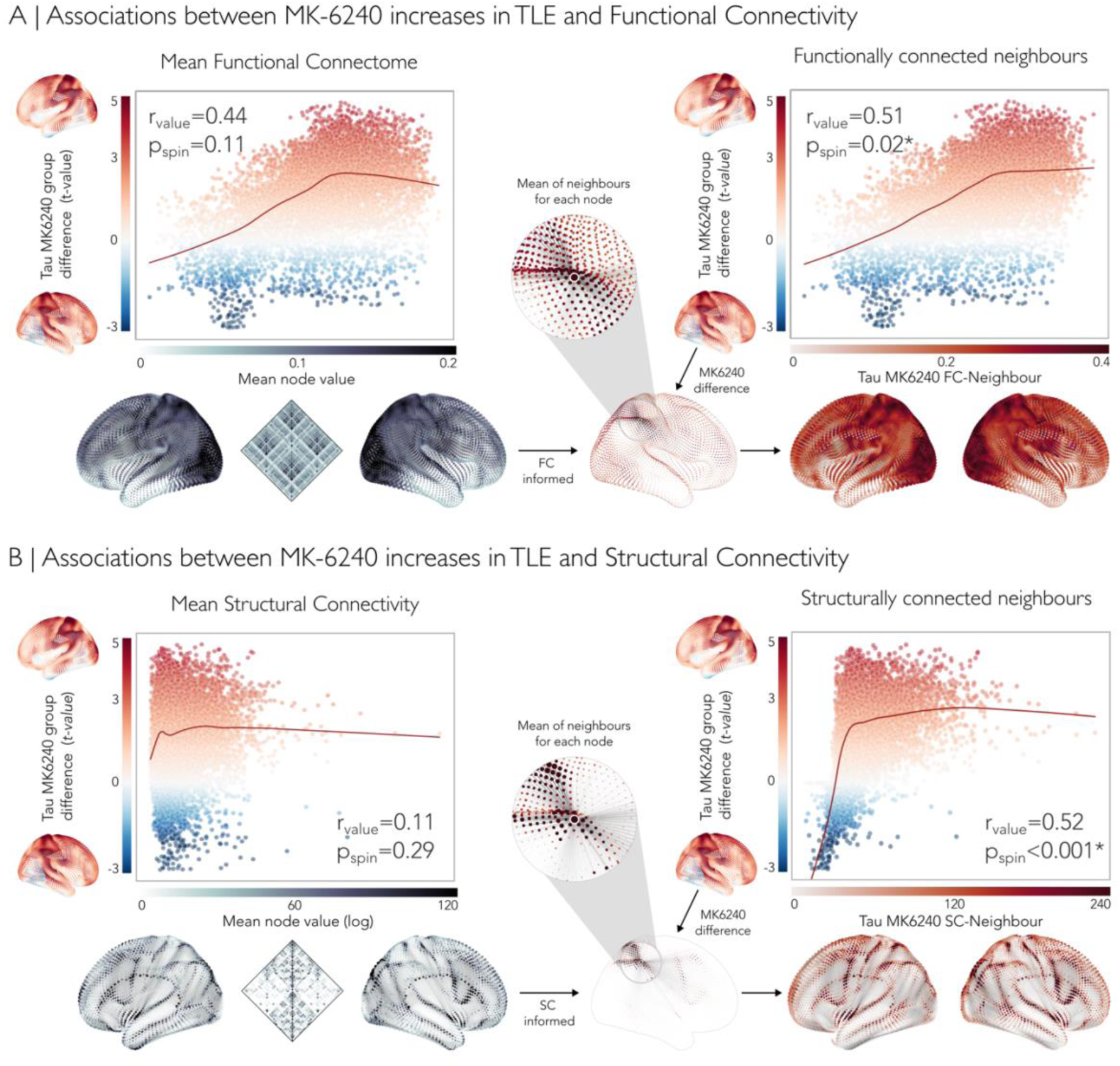
**A)** Relationship between MK-6240 group differences and functional connectivity. The left panel displays a scatterplot of mean node values from the functional connectome against between-group differences in MK-6240 tau uptake (see Figure 1). The right panel visualizes neighboring regions of each node determined by functional connectivity, with their association to tau between-group differences. **B)** Relationship between MK-6240 group differences and structural connectivity, derived from diffusion MRI tractography. The left panel shows a logarithmic scatter plot of mean node values against group differences. The right panel highlights neighboring regions based on structural connectivity, which show a strong trend of increased tau deposition. The central image depicts how functional (upper) and structural (lower) connectivity informed by MK-6240 group differences was used to generate neighbor-weighted MK-6240 estimates. Points are color-coded based on MK-6240 t-values, with higher t-values indicating greater tau deposits in TLE compared to controls. The red line represents non-linear trends in the data. These findings suggest that both functional and structural network properties significantly reflected the spatial distribution of MK-6240 increases in TLE, with network hubs and their connected regions being particularly vulnerable.

This analysis revealed a marginal association at trend levels between MK-6240 SUVR and highly connected regions defined by mean functional connectivity (r = 0.44, p_spin_ = 0.11, **Figure 3A-left**). Associations with mean structural connectivity were non-significant (r = 0.11, p_spin_ = 0.29, **Figure 3B-left**).

In contrast, associations between tau uptake differences and neighborhood-weighted connectivity showed strong correlations for both functional (r = 0.51, p_spin_ 0.02, **Figure 3A-right**) and structural networks (r = 0.52, p_spin_ < 0.001, **Figure 3B-right**).

### Direct effect of Tau on memory surpasses network-mediated influence

In a final integrative analysis, we evaluated the effects of MK-6240 on cognitive performance and the potential modulation of the structural connectome using structural equation modeling (SEM). In the SEM analysis (**Figure 4),** mean MK-6240 SUVR from significant clusters was significantly associated with structural neighborhood-weighted connectivity (β = 0.38, p = 0.002), but not with structural connectome efficiency. Higher MK-6240 was directly and significantly associated with lower EpiTrack performance (β = −0.49, p = 0.009) and lower episodic memory performance (β = −0.55, p = 0.004), but not with semantic memory (β = −0.14, p = 0.56). Paths from structural network measures to memory outcomes were not significant, and none of the indirect (mediation) effects reached significance. The total effect of MK-6240 on episodic memory was significant, indicating a predominantly direct relationship rather than mediation.

**Figure 4.**
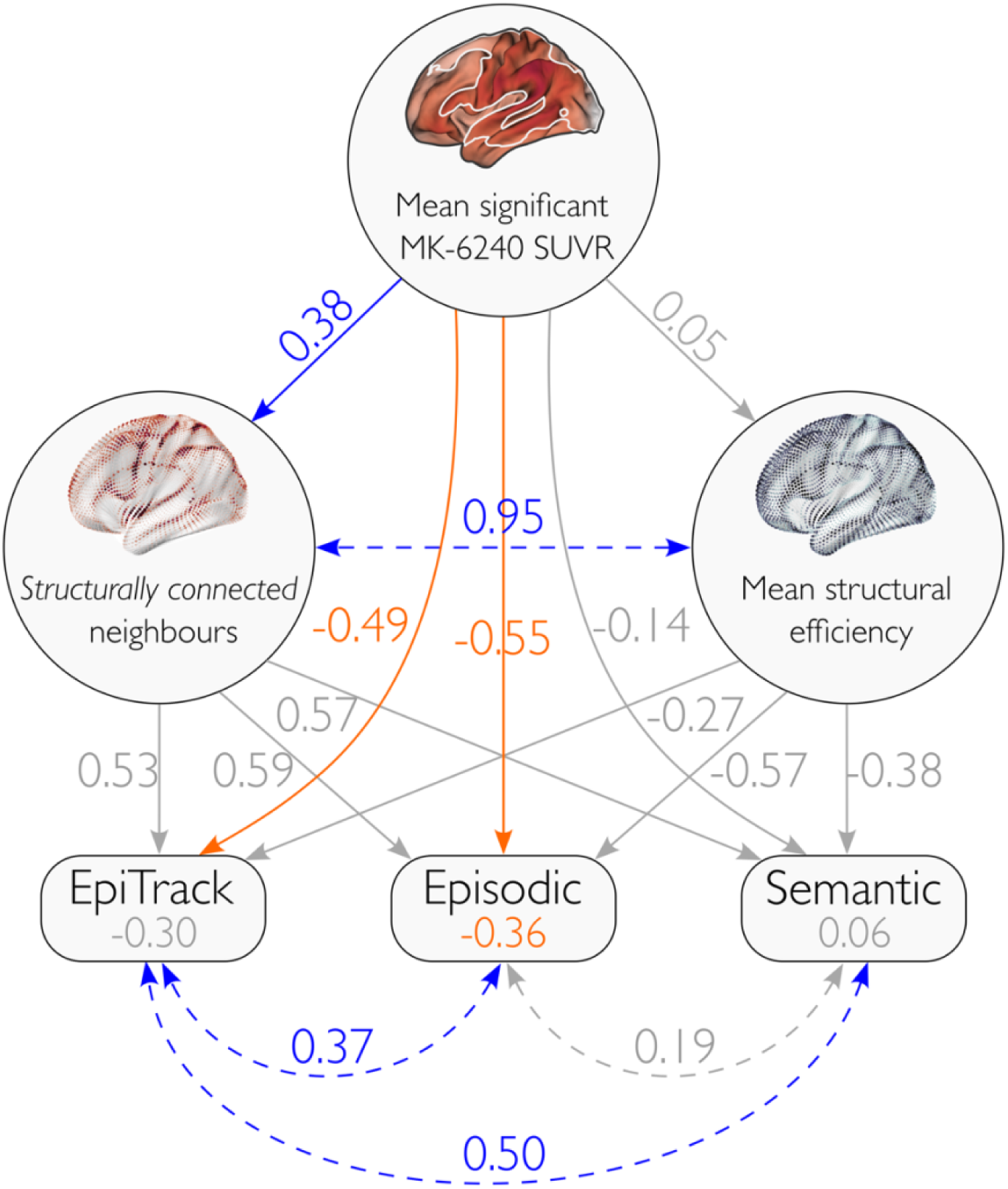
Structural equation model showing relationships between mean significant MK-6240, structural network measures (SC weighted neighbors and SC efficiency), and behavioral outcomes. Values represent standardized path coefficients. Significant paths (p < 0.05) are shown in blue for positive effects and orange for negative effects, the rest is gray. Dotted lines represent covariance between features. Indirect effects via semantic network measures were tested but were not significant.

### Heterogeneous AT8 Tau depositions in post-surgical TLE tissue

Neuropathological evaluation of AT8-immunoreactive phosphorylated tau (p-tau) revealed a spectrum of variable p-tau deposition in 5/6 cases for which adequate specimen was available (**Supplementary Table 5**). The most frequent finding was focal subpial immunoreactivity, characterized by a thin layer of positivity at the cortical margin, which was particularly prominent in cases diagnosed with focal cortical dysplasia (FCD Type II and III). Intracytoplasmic neuronal inclusions were identified in several instances, most notably in a case with a hippocampal cavernoma, where perilesional neurons exhibited definite intracytoplasmic staining alongside strong “tuffed-like” pathology in adjacent areas. In a case of FCD-II the morphology was more complex, presenting as strong neuritic staining within the hippocampal CA4 region and focal, strong longitudinal striated patterns. One case had less suitable specimen, but overall exhibited negative findings, with only negligible dot-like staining at fragment edges (**Supplementary Figure 9**).

## Discussion

This study represents novel *in-vivo* evidence of elevated phosphorylated tau, measured by [¹⁸F]MK-6240 PET, in young adults with TLE, and demonstrates association with markers of network architecture as well as clinical and cognitive measures. Comparing patients to controls, increased tau accumulation in TLE was observed bilateral temporo-posterior regions, and we could furthermore demonstrate an association with measures of functional and structural connectivity. The latter findings indicate that highly interconnected network hubs and their neighboring regions are particularly vulnerable to tau deposition. Notably, females with TLE exhibited a higher tau burden than males, suggesting potential sex-related differences in tau pathology. Cognitive analyses showed that higher tau burden was directly associated with lower performance on the Epitrack and impaired episodic memory. Mediation analyses did not support indirect effects of structural network measures on cognitive outcomes, indicating that the relationship between tau burden and memory impairment was predominantly direct rather than mediated by network properties. Although longitudinal changes were modest, these findings suggest that tau pathology may contribute to disease-related network disruption and cognitive impairment in drug-resistant TLE. These findings suggest that tau accumulation may contribute to cognitive decline, supporting the potential role of tau in neurodegeneration in pharmaco-resistant epilepsy *in-vivo*.

While so far unexplored in epilepsy research, the [^18^F]MK-6240 PET tracer has been repeatedly applied to map disease processes in established neurodegenerative conditions, in particular AD^28–30^. In those assessments, MK-6240 has been shown to generally describe the typical Braak pattern of pathological spread away from an entorhinal/mesiotemporal disease epicentre to involve bilateral, large scale networks^31^. Several assessments in AD have also shown that tau PET changes have associations with overall cognitive impairment and atrophy, both at the time of scanning^32,33^ and prospectively^34^. These findings have been complemented with several histopathological and electron microscopic validation experiments^35^, suggesting that MK-6240 does scale with the overall deposition of hyperphosphorylated tau. A key strength of our study is its focus on young and middle-aged individuals, an underexplored population in tau imaging research. Despite the younger age of the cohort, we observed a robust increase in MK-6240 binding in our patients. The observed pattern of tau deposition, involving the superior and middle temporal regions, as well as parietal-occipital areas, aligns in part with earlier *ex-vivo* observations of increased abnormalities in the ipsilateral temporal lobe relative to the epileptogenic focus^3,10,36–38^, and confirms recent CSF-biomarker findings suggesting increased tau in TLE^39^. These results provide *in-vivo* evidence suggesting that TLE is associated with an elevated risk of tau-associated pathophysiological processes^39,40^. Crucially, increases in tau deposition in TLE patients relative to controls remained after adjusting for age and sex, with findings remaining significant after correction for multiple comparisons. Prevalence mapping provided evidence for slightly greater tau burden ipsilateral to the seizure focus, whereas group-level effects were largely bilateral. This pattern was not directly aligned with ipsilateral temporo-limbic connectivity and may therefore reflect a more complex pathological process that cannot be fully explained with respect to a unilateral mesiotemporal epicentre.

Our sample was composed out of equal proportions of females and males in both the patient and control cohorts. This allowed us to examine interactions between group and sex, which pointed towards more marked between-group differences in females compared to males. Indeed, while both female and male patients demonstrated common increases in tau uptake in central and posterior divisions, female patients additionally displayed increases in tau compared to female controls in anterior midline, insular, and temporo-anterior divisions. In the context of patients diagnosed with AD and individuals with a familiar risk for it, sex effects have increasingly been recognized. This is also motivated by a twofold higher lifetime risk for developing AD in women, which cannot be explained by higher life expectancy alone^41^. Prior studies have indicated higher tau accumulation in both the brain and plasma in older females compared to males that are both cognitively normal and those that are at risk for developing AD. Increased tau in females is generally not accompanied by an elevated increase in amyloid, suggesting either independent mechanisms or an elevated effect of pre-existing amyloid on tau accumulation in females^42,43^. While the evidence for sex-differences in longitudinal tau effects remains inconclusive, there is work that overall suggests faster cognitive decline in females with significant tau accumulation compared to their male counterparts^44^. It has been suggested that sex hormones and sex chromosomes interact with various disease mechanisms during aging, affecting inflammatory, metabolic, and autophagy processes^45^. These complex interactions affect disease progression between the sexes in classical neurodegenerative conditions and may furthermore interact with disease related factors and reported sex differences in epilepsy^46^.

Previous PET studies in TLE have primarily focused on functional rather than pathological markers, with FDG-PET commonly used to map regional hypometabolism for seizure lateralization, surgical planning, and outcome prediction^52–54^. Cross-sectional studies have reported moderate associations between epilepsy duration and FDG-PET hypometabolism, mainly in temporal lobe regions^55,56^. In contrast, longitudinal FDG-PET evidence remains limited and inconclusive. A study of 16 adults with TLE undergoing repeated FDG-PET scans found no significant association between scan interval and the rate of change in regional FDG-PET uptake, although patients with longer disease duration showed greater evidence of progressive decline^57^. Similar findings have been reported in pediatric non-lesional epilepsy, where persistent seizures were associated with expansion of the hypometabolic cortex over time, whereas seizure reduction was linked to decreased hypometabolic extent^58^.

Connectome decoding models have previously been applied to describe network effects in the expression of specific imaging phenotypes, in both neurodevelopmental and neurodegenerative conditions^20,59–62^. In our study, we explored global as well as local connectivity associations to patient-related MK-6240 increases. While the former approach aimed to test an overall susceptibility of highly interconnected hub regions to tauopathy, the latter assessed a more local association between tau uptake and a region’s connectivity patterns. Notably, while TLE-related increases in tau showed only marginal associations with global functional connectivity, stronger associations emerged with local connectivity patterns, both structurally and functionally. These findings suggest a subtle vulnerability of polysynaptic functional hubs, alongside a more pronounced influence of local network architecture. The selective susceptibility of highly connected cortical hubs in TLE has been described in studies of atrophy, and our findings extend this vulnerability to tau pathology^63^. In neurodegeneration research, local connectivity has been linked to the mechanisms of regional propagation, in which tau spreads through anatomically connected networks to affect an increasing set of brain regions^31,64,65^.

In clusters of between-group differences, we did not find an association between MK-6240 and either disease duration or age. However, surface-based analyses revealed that longer disease duration was associated with increased MK-6240 SUVR in contralateral parieto-occipital regions, while age was associated with increased SUVR in anterior ipsilateral inferior cingulate and inferior frontal regions. These distributed effects may reflect a broader disease process involving interconnected networks rather than localized effects limited to the ipsilateral temporal lobe^14^. Increasing the sample size with longer longitudinal follow-up times and an older average age of included patients may boost sensitivity to detect progressive tau uptake in TLE in future work, and might help to clarify the temporal dynamics of tau accumulation and its potential role in neurodegenerative cascades. This expanded design help to identify clinically relevant subgroups that show the most marked progressive increases in tau, and to establish whether tau increases are pre-existing or whether they cumulatively increase throughout the disease course in TLE. In AD, prior work has reported an increase in misfolded tau deposition over periods longer than five years and in individuals older than 65 years^66,67^. We did not systematically genotype our participants to explore potential associations between *in-vivo* tau uptake and genetic background. A prior meta-analysis of 46 studies^68^ found that the APOE ε4 allele^69^ was associated with increased epilepsy risk, earlier onset, worse cognition, and a history of febrile convulsions, while the APOE ε2 allele was linked to reduced risk; APOE ε3 showed no association^70^. The APOE gene was more strongly related to temporal lobe, drug-refractory, and late-onset epilepsy, with no link between ε4 and psychiatric symptoms or seizure-free outcomes after surgery^8^.

In AD, increased tau burden is associated with reduced hippocampal volume, likely reflecting neuronal loss driven by abnormal tau deposition and network disconnection from memory systems^71–73^. Hippocampal atrophy has also been linked to high amyloid-β burden in tau-positive cases and to cognitive decline^74^. In our study, we observed an association between increased cortical MK-6240 uptake and reduced ipsilateral hippocampal volume. This relationship may reflect network-level effects that impact the hippocampus together with large-scale systems. In several prior studies, associations between hippocampal pathology and network level compromise have been demonstrated^75–77^, suggesting a coupling of pathological processes in TLE occurring the mesiotemporal lobe and at large scale. However, detailed analyses of these mechanisms beyond the correlational assessment conducted here are beyond the scope of the present manuscript, and further studies are needed to clarify the role of hippocampal tau pathology in cognition and network disconnection.

Our findings also demonstrated significant associations between MK-6240 uptake and poorer performance in executive function and episodic memory, which again support a potential role of phosphorylated tau in progressive, multisystem dysfunction in TLE. These findings suggest a vulnerability of cognitive networks governing episodic memory and executive function to tau-related pathology in TLE. The preservation of semantic memory in our patients and the absence of a correlation with MK-6240 may, on the other hand, suggest compensatory mechanisms similar to those in AD, where episodic memory and executive function are more markedly affected than semantic memory and conceptual knowledge^3,78^. In future work, the incorporation of other measures probing into additional mechanisms of memory dysfunctions, as well as those tapping into processes mediated by other systems (*e.g.,* parieto-frontal control processes) may be valuable.

Post-surgical neuropathological analysis using the AT8 antibody revealed p-tau deposition in 5/6 cases with available specimen, providing histopathological support for our *in-vivo* PET findings. Several studies have shown that p-tau accumulation correlates with duration of refractory epilepsy and cognitive decline^10,14,36,38^, appearing across a diverse spectrum of surgical aetiologies including FCD, hippocampal sclerosis, and acquired lesions^47^. Our findings align with these observations. On the other hand, the variability of tau staining across specimens underscores the complexity of tau pathology in epilepsy. While the PET signal most likely reflects overall increases in tau burden, findings may also co-localize with inflammatory cascades, given reports linking cortical atrophy patterns to microglial activation in TLE^48,49^. Growing evidence further suggests that neuroinflammation may accelerate tau phosphorylation and misfolding, creating a self-perpetuating cycle of neuronal dysfunction^50,51^.

Together, these findings suggest that elevated phosphorylated tau in young adults with TLE may represent a candidate mechanism contributing to large-scale network dysfunction. As tau imaging in epilepsy advances, integrating larger, longitudinal, and multimodal datasets across diverse age ranges will be critical to clarifying disease trajectories and individual variability. Ultimately, this work lays the groundwork for a network-based molecular framework of epilepsy, which can be refined as more high-resolution *in-vivo* and *post-mortem* data become available.

## Methods

### Participants

We recruited 28 TLE patients (14 Females, mean±SD age=33±8 years) and 28 healthy volunteers with a similar age and sex distribution (14 Females, mean±SD age=37±12 years) between October 2019 and July 2025. Diagnosis of TLE and lateralization of seizure foci were determined by the criteria of the International League Against Epilepsy (ILAE), based on a comprehensive examination that included clinical history, neurological examination, review of medical records, video-electroencephalography (EEG) recordings, and clinical MRI. As in earlier work^75^, hippocampal atrophy was determined as volumes or left-right asymmetry beyond 1.5 standard deviations of the corresponding mean of healthy controls: 8 (29%) patients had unilateral hippocampal atrophy ipsilateral to the focus and 19 (71%) had normal hippocampal volume. Mean±SD age at seizure onset was 21±14 years and mean±SD duration of epilepsy was 15±11 years. Healthy controls were recruited via advertisement; all denied a history of neurological, psychiatric, or significant medical comorbidities, including traumatic brain injury. A subgroup in both cohorts had longitudinal acquisition (28/13 patients and 28/7 controls) with a minimum follow up time of 12 months (median=15 months, range=12 months – 42 months). Interictal epileptiform discharge frequency (IEDs) was obtained either from the epilepsy monitoring unit or from the long-term monitoring (EEG). Generalized tonic-clonic seizures frequency (GTCS) was obtained through retrospective clinical data review.

### Standard protocol approvals, registrations, and patient consents

The Ethics Committee of the Montreal Neurological Institute and Hospital (MNI) approved the study (#2018-4148). Written informed consent, including a statement for open sharing of collected research data, was obtained from all participants.

### PET Image acquisition and processing

Tau PET scanning was performed on a Siemens high-resolution research tomograph (HRRT) at the McConnell Brain Imaging Center of the MNI, using [^18^F]-MK-6240 tracer^17^, which shows low off-target binding^16^. The tracer was injected intravenously, and images were acquired after 90 minutes for approximately 30 minutes. A previous validation study in our center has shown that scans 90-110 minutes post-injection may optimally balance duration and signal. In TLE, mean injected MK-6240 activity was 226.1±22.6 MBq/L. In controls, it was 237.2±21.5 MBq/L. Images were reconstructed with an ordered subset expectation maximization algorithm. A short transmission scan (11 minutes) with a rotating Cesium137 source was used for attenuation correction. Images underwent correction for dead time, decay, as well as random and scattered coincidences. Dynamic MK-6240 images (4 in total) were co-registered and averaged. Average MK-6240 uptake was registered to the T1-weighted MRI (T1w) using a novel affine, edge-based approach with a cross-correlation cost function and histogram matching (**Supplementary Figure 7A**). Next, it was partial volume corrected using a probabilistic mask of the grey matter. We calculated standardized uptake value ratio (SUVR) images using the cerebellar grey matter as a reference region^79^. To assess reproducibility in the mixed effects models, we additionally tested the use of the brainstem and a composite reference region^80^ (**Supplementary Figure 3**). The SUVR volume was registered to the medial cortical surface derived from T1-weighted MRI, resulting in surface-based MK-6240 maps (**Supplementary Figure 7B**). Surface-based MK-6240 data were blurred with a 10-mm FWHM surface-based kernel, and participant-specific data were re-sampled to the fsLR-32k standard template surface^81^. The following subcortical structures were automatically segmented on the T1w image for each hemisphere: thalamus, caudate nucleus, putamen, globus pallidus, nucleus accumbens, as well as the hippocampus After calculating the mean value of all the voxels of each structure, we computed SUVR values based on the same reference region as for the other structures.

### Cognitive assessments

Cognitive assessment included the EpiTrack, a 15-min screening tool primarily targeting evaluate executive function, designed to monitor cognitive side effects of anti-seizure medication and seizures^79^. Episodic and semantic memory were assessed using previously validated tasks. While the episodic memory task involved learning/recalling object pairs across encoding and retrieval phases, the semantic memory task required identifying conceptually related object pairs from visual prompts^80,81^. For logistic reasons, the cognitive tests were performed at the time of the MRI scanning (13±15 months apart from the PET scanning across participants).

### Multimodal MRI acquisition and processing

MRI scans were also obtained at the McConnell Brain Imaging Centre, using a 3T Siemens Magnetom Prisma-Fit scanner equipped with a 64-channel head coil. Participants underwent a T1w structural scan, followed by multi-shell diffusion weighted imaging (DWI) and resting-state functional MRI (rs-fMRI). T1w images were acquired using a 3D-MPRAGE sequence (0.8 mm isovoxels). Diffusion-weighted imaging data were collected using a multi-band accelerated 2D spin-echo EPI sequence (1.6 mm isovoxels) with three b-value shells (300, 700, 2000 s/mm²) and corresponding diffusion directions (10, 40, 90), along with phase-reversed b0 images for distortion correction. The rs-fMRI scans were acquired with a 7-min multiband 2D-BOLD EPI sequence (3 mm isovoxels, TR = 600 ms), with participants instructed to keep their eyes open and fixate on a cross. For detailed parameters, see the MICs database (http://n2t.net/ark:/70798/d72xnk2wd397j190qv)^82^.

Multimodal MRI data processing and feature fusion leveraged micapipe v0.2.3 (http://micapipe.readthedocs.io), with details described elsewhere^83^. In brief, T1w data were processed for intensity abnormalities, followed by automated segmentation of subcortical structures using FSL FIRST^84^, the generation of probabilistic tissue estimates using FSL FAST^85^, and cortical surface segmentations using FreeSurfer 6.0^86^. Surfaces were manually quality controlled and corrected if necessary.

The rs-fMRI data underwent motion correction, distortion correction, denoising, bandpass filtering, surface mapping, and surface registration. We computed pairwise Product-moment correlation coefficients between vertices to generate a subject-level functional connectivity (FC) matrix. Resulting correlation coefficients were normalized via Fisher r-to-z transformation.

DWI preprocessing involved motion correction, denoising, and correction for inhomogeneity and geometric distortions. Anatomically constrained tractography was performed using tissue segmentations derived from each participant’s T1w images that were co-registered to native DWI space. Multi-shell and multi-tissue response functions were estimated, followed by constrained spherical deconvolution and intensity normalization. A tractogram with 40M streamlines was generated and refined using SIFT2^87^. Structural connectome (SC) weights were defined as the filtered streamline counts between cortical vertices using the fsLR-5k standard surface. A secondary SC was generated using streamline edge lengths as weights. These two connectomes were combined to create a weighted SC (SCw) that integrates connectivity strength and distance information, such that the connection with high streamline counts but short lengths are penalized relative to longer connections.

### Data Normalization

A novel PET-to-MRI registration was performed in two steps. First, edge images were generated from the T1w and the average MK-6240 PET map. Next, the edge image of the MK-6240 average map was registered to the T1w edge image using a cross-correlation cost function with histogram matching. (**Supplementary Figure 7A**). Individual quality control was performed for each registration to ensure optimal results. For DWI and rs-fMRI, a two-step registration process was applied. First, a modality-agnostic, deep learning-based segmentation^88^ was used to generate labeled images for each modality. These labeled images were then registered to the T1w-derived label map using affine registration, followed by nonlinear registration to ensure alignment in the subject’s native T1w space.

The resulting transformations were applied to map PET data onto the subject’s cortical surface, reconstructed from the native T1w image. This native surface was subsequently resampled to the fsLR standard surface meshes at two different resolutions ∼32,000 and ∼5,000 vertices per hemisphere. The fsLR-32k surface was used to sample quantitative values from MK-6240, while the fsLR-5k surface was used for constructing both structural and functional connectomes derived from DWI and rs-fMRI.

### Statistical analysis

Analyses were performed using R and BrainStat^89^, an open-access toolbox for brain-wide statistical analysis (http://brainstat.readthedocs.io). In patients, prior to analysis, cortical features were sorted into ipsilateral/contralateral relative to the seizure focus.

*a) Mapping of tau uptake in TLE.* We compared MK-6240 vertex-wise SUVR between individuals with TLE and healthy controls using a surface-based linear mixed effects model, controlling for age, sex, and participant as a random effect. Separate surface-based models were applied to control for cortical thickness and to compare males with TLE versus male controls, and females with TLE versus female controls. An additional interaction model was used to assess the combined effects of group and sex on tau uptake. Here and below, surface-based analyses were corrected for multiple comparisons using random field theory (two-tailed), with a significance threshold set at p < 0.025. As in the cortical analysis, we compared MK-6240 uptake in subcortical ROI between TLE patients and healthy controls using mixed effects models. Models included age as a covariate, a sex by group interaction, and a random intercept for each participant. Statistical significance was assessed using a two-tailed threshold of p < 0.025, with false discovery rate correction for multiple comparisons.

In addition to examining between-group differences, a binary tau probability map was created by thresholding the MK-6240 PET data in every individual. The threshold was defined as the 97.5th percentile (1.56 SUVR) of values in the healthy control population (**Table 2**). Vertices exceeding this threshold were assigned a value of 1, while all other vertices were set to 0, resulting in a binary map indicating regions of elevated tau deposition. The group difference in mean MK-6240 SUVR values from significant clusters (MK-6240_sig_) in both hemispheres was assessed using a Welch two sample t-test.

Longitudinal changes in MK-6240 were assessed using linear-mixed-effects models. One model evaluated the change in MK-6240 uptake over time from the mean values extracted from significant regions. This model was fitted using restricted maximum likelihood (REML), and t-tests for fixed effects were calculated using Satterthwaite’s approximation for degrees of freedom. Separate estimated slopes and 95% confidence intervals were obtained for each group using the Kenward-Roger method. Model assumptions were checked via inspection of scaled residuals, and between-subject variability was captured by the participant-level random intercept. A second model evaluated surface-based longitudinal changes in MK-6240 uptake comparing TLE and healthy controls. This model was adjusted for age and sex and corrected for multiple comparisons using random field theory (p<0.025).

*b) Relationship to clinical and cognitive variables.* Correlation analyses were initially performed using mean MK-6240 SUVR values extracted from significant clusters in both hemispheres identified in the group-difference analysis (**Figure 2, left column**). These findings were complemented with surface-based linear models using vertex-wise MK-6240 SUVR values (Figure 2, right column). These analyses included only cross-sectional data from the first time point. We examined associations to age, disease duration, age at onset, ipsilateral/contralateral hippocampal volume, and cognitive measures. Unless otherwise specified, these analyses controlled for age and sex. In addition to the analyses of Figure 2 we evaluated the relationships between MK-6240 and age at onset of the epilepsy and the contralateral hippocampal volume, but no significant correlations were observed (**Supplementary Figure 5**).

*c) Network contextualization.* We correlated the t-statical map from the above between-group comparison in MK-6240 between patients and controls with network metrics derived from group-level SC and FC. Specifically, network metrics were mean connectivity and neighborhood-weighted connectivity^90^. SCs and FCs were modeled as non-thresholded, weighted graphs. Each subject’s connectome was represented by a vertex-wise adjacency matrix based on either structural or functional connectivity. Group-level connectomes were then constructed by averaging edge weights across individuals, separately for DWI and rs-fMRI data.

From the group-level weighted connectivity graphs, two primary network measures were derived: mean connectivity and neighborhood-weighted connectivity, the latter also referred to as Neighborhood Deformation Estimates^90^. These metrics were used as dependent variables in the group-level correlation analyses, contextualizing regional differences in MK-6240 findings (t-statistical map) within the broader network architecture.

To compute neighborhood-weighted connectivity, we applied the following procedure. For each cortical node (*D_i_*), the feature values of its connected neighbors were extracted and averaged (*dj*), weighted by the strength of their connection (*NC_ij_*). Self-connections (i.e., diagonal elements of the connectivity matrix, *j≠i*) and the medial wall vertices were excluded from the computation to avoid circular influence. The neighborhood-weighted value for a given node was thus a degree-normalized, connectivity-weighted average of its neighbors’ feature values, such as the t-statistical map of MK-6240 difference. See **Equation 1**:

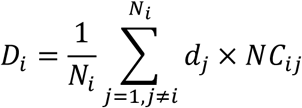

This method quantifies the relation between the original nodal measurements, for example regional PET uptake, and the connectivity-weighted average of these same measurements across each node’s network-defined neighborhood. In doing so, it captures the extent to which regional signal patterns are spatially structured by the underlying network topology. Detailed implementation and supporting scripts are available in our project repository: https://github.com/MICA-MNI/2025_in-vivo_tauPET-mk6240_TLE/blob/main/scripts/Fig-2_network_contextualization.ipynb

To correct for multiple comparisons and account for the spatial autocorrelation inherent in surface-based data, non-parametric spin permutation testing was used (using 10,000 permutations)^91,92^. Significance was assessed relative to these null models. Medial wall vertices were excluded from all calculations.

*d) Structural equation modeling.* SEM examined whether graph-theoretical measures of structural brain networks (see below) were associated with the relationship between MK-6240sig SUVR and three memory outcomes (EpiTrack, episodic memory, and semantic memory). Models were estimated using maximum likelihood with non-parametric bootstrapping (10,000 resamples) to obtain robust standard errors and confidence intervals, particularly for indirect effects. Model fit was summarized using the root mean square error of approximation and the standardized root mean square residual, and standardized path coefficients are reported.

Graph-theoretical metrics were derived from individual fsLR-5k structural and functional weighted connectomes^93^. For each participant, we computed mean strength, mean neighborhood-weighted connectivity, mean characteristic path length, weighted global efficiency^94^, and weighted clustering coefficient^95,96^. Functional connectomes were Fisher-z transformed, absolute values were taken, normalized to the maximum weight, and thresholded to retain the top 20% of connections^97,98^. Structural connectomes were weighted by edge length to penalize short-distance connections with high connectivity and subsequently normalized to the maximum weight. Individual mean neighborhood-weighted connectivity was calculated using each subject’s weighted structural connectome (*NC_ij_*) with MK-6240 uptake values from connected cortical neighbors (*di*, see **Equation 1**). The medial wall was removed from all connectomes. To explore associations among variables, correlograms were generated including MK-6240sig, behavioral measures, and network metrics. Due to high collinearity among network measures, only structural weighted global efficiency and structural mean neighborhood-weighted connectivity were retained for SEM analyses. Functional network metrics were excluded because they showed no significant associations with behavioral outcomes (**Supplementary Figure 8**).

Three preliminary SEMs were first estimated separately for each cognitive outcome (EpiTrack, episodic memory, semantic memory). Based on these results, a single multivariate SEM was specified with MK-6240sig as the predictor, structural weighted global efficiency, and mean neighborhood-weighted connectivity as mediators, and the three memory measures as correlated outcomes. The predictor was specified to directly predict both mediators (paths *a1* and *a2*), which were allowed to covary. Each outcome was then regressed on both mediators and on MK-6240 signal, estimating mediator-to-outcome (*b*) paths and direct (*c′*) effects. Residual covariances were specified among outcomes to account for shared unexplained variance. Indirect effects were calculated as the product of *a* and *b* paths for each mediator and outcome, and total indirect effects were obtained by summing mediator-specific indirect effects. Total effects were computed as the sum of direct and indirect effects. All parameters were estimated simultaneously, and full model estimates are reported in **Table 4**.

**Table 4.** Main Structural Equation Model Summary.

| Regressions | Estimate | Std.Err | z-value | P(>\|z\|) | Std.lv | Std.all |
| --- | --- | --- | --- | --- | --- | --- |
| *SC neighbours ~*<br>***MK-6240 (a1)*** | **0.17** | **0.056** | **3.051** | **0.002** | **0.17** | **0.378** |
| SC efficiency ~<br>MK-6240 (a2) | 0.053 | 0.164 | 0.321 | 0.748 | 0.053 | 0.045 |
| EpiTrack ~ |  |  |  |  |  |  |
| SC neighbours (b11) | 9.945 | 8.821 | 1.127 | 0.26 | 9.945 | 0.534 |
| SC efficiency (b12) | -1.92 | 3.31 | -0.58 | 0.562 | -1.92 | -0.267 |
| ***MK-6240 (c1)*** | **-4.08** | **1.557** | **-2.62** | **0.009** | **-4.08** | **-0.486** |
| Episodic ~ |  |  |  |  |  |  |
| SC neighbours (b21) | 9.769 | 9.801 | 0.997 | 0.319 | 9.769 | 0.586 |
| SC efficiency (b22) | -3.656 | 3.481 | -1.05 | 0.294 | -3.656 | -0.567 |
| ***MK-6240 (c2)*** | **-4.148** | **1.456** | **-2.848** | **0.004** | **-4.148** | **-0.552** |
| Semantic ~ |  |  |  |  |  |  |
| SC neighbours (b31) | 8.348 | 6.701 | 1.246 | 0.213 | 8.348 | 0.567 |
| SC efficiency (b32) | -2.158 | 2.075 | -1.04 | 0.298 | -2.158 | -0.379 |
| MK-6240 (c3) | -0.921 | 1.559 | -0.591 | 0.555 | -0.921 | -0.139 |
| Covariances | Estimate | Std.Err | z-value | P(>\|z\|) | Std.lv | Std.all |
| *SC neighbours ~*<br>*SC efficiency* | **0.011** | **0.002** | **6.041** | **>0.001** | **0.011** | **0.954** |
| EpiTrack ~ |  |  |  |  |  |  |
| *Episodic* | **0.457** | **0.174** | **2.624** | **0.009** | **0.457** | **0.373** |
| *Semantic* | **0.577** | **0.194** | **2.979** | **0.003** | **0.577** | **0.501** |
| Episodic ~ |  |  |  |  |  |  |
| Semantic | 0.194 | 0.134 | 1.446 | 0.148 | 0.194 | 0.189 |
| Variances | Estimate | Std.Err | z-value | P(>\|z\|) | Std.lv | Std.all |
| SC neighbours | 0.004 | 0.001 | 6.145 | >0.001 | 0.004 | 0.857 |
| SC efficiency | 0.031 | 0.005 | 6.029 | >0.001 | 0.031 | 0.998 |
| EpiTrack | 1.37 | 0.337 | 4.065 | >0.001 | 1.37 | 0.848 |
| Episodic | 1.094 | 0.173 | 6.328 | >0.001 | 1.094 | 0.844 |
| Semantic | 0.969 | 0.4 | 2.425 | 0.015 | 0.969 | 0.957 |
| Regressions | Estimate | Std.Err | z-value | P(>\|z\|) | Std.lv | Std.all |
| Indirect EpiTrack > SC<br>neighbours | 1.694 | 1.595 | 1.062 | 0.288 | 1.694 | 0.202 |
| Indirect EpiTrack > SC<br>efficiency | -0.101 | 0.633 | -0.16 | 0.873 | -0.101 | -0.012 |
| Indirect EpiTrack > Total | 1.593 | 1.312 | 1.214 | 0.225 | 1.593 | 0.19 |
| Indirect Episodic > SC<br>neighbours | 1.664 | 1.844 | 0.903 | 0.367 | 1.664 | 0.221 |
| Indirect Episodic > SC<br>efficiency | -0.193 | 0.894 | -0.216 | 0.829 | -0.193 | -0.026 |
| Indirect Episodic > Total | 1.471 | 1.505 | 0.978 | 0.328 | 1.471 | 0.196 |
| Indirect Semantic > SC<br>neighbours | 1.422 | 1.202 | 1.183 | 0.237 | 1.422 | 0.214 |
| Indirect Semantic > SC<br>efficiency | -0.114 | 0.526 | -0.216 | 0.829 | -0.114 | -0.017 |
| Indirect Semantic > Total | 1.308 | 1.003 | 1.305 | 0.192 | 1.308 | 0.197 |
| Total EpiTrack | -2.487 | 1.339 | -1.857 | 0.063 | -2.487 | -0.296 |
| **Total Episodic** | **-2.677** | **0.988** | **-2.71** | **0.007** | **-2.677** | **-0.356** |
| Total Semantic | 0.388 | 1.166 | 0.332 | 0.74 | 0.388 | 0.058 |

**Table 5.** Post-surgical TLE cases.

| ID | Surgical Procedure | Histology | AT8+ | Neuropathology |
| --- | --- | --- | --- | --- |
| S1 | Right CAH | FCD-II/HS | yes | Focal subpial and few intracytoplasmic neuronal inclusions. Weak subpial staining with negative neurons. |
| S2 | Right AH | FCD-IIA/HS | yes | Subpial positivity and rare isolated neurons with weak staining. |
| S3 | Right fusiform and parahippocampal resection | FCD-? | no | Neurons negative; dot-like edge staining only. |
| S4 | Left CAH | FCD-IIA | yes | Weak subpial stain and rare neuronal positivity. Otherwise, negative with focal background |
| S5 | Right CAH | FCD-IIIA/HS2 | yes | Weak subpial staining is present, with a few neurons showing intracytoplasmic positivity, alongside strong neuritic staining in CA4 and focal longitudinal striated staining. |
| S6 | Left amygdalar resection | Cavernoma | yes | Definite intracytoplasmic neuronal positivity around cavernoma; strong tufted-like staining in focal areas |

2 *CAH: cortico amygdalo hippocampectomy*3 *AH: amygdalo hippocampectomy*

### Tau immunohistochemistry

Brain tissue samples from the surgical resections of the temporal neocortex, parahippocampal regions, hippocampus, and amygdala were processed according to standardized clinical procedures at the Montreal Neurological Institute. Samples were fixed in 10% neutral buffered formalin and embedded in paraffin (FFPE). Immunohistochemistry was carried on a Ventana Discovery Ultra automated staining platform, following manufacturer instructions. The sections were pre-treated with CC1, and the primary monoclonal antibody AT8 (recognizing phosphorylated tau at Ser202/Thr205; Thermo Fisher Scientific; MN1020) used at 1:1200 dilution. Whole-slide imaging was performed using a Hamamatsu NanoZoomer S210 digital scanner (C13239-01). All digitized images were reviewed and interpreted by an expert neuropathologist to characterize the distribution and morphology of phosphorylated tau inclusions.

## Data availability

All the data used in the study is hosted on OSF (https://osf.io/ct3gw).

## Code availability

All processing workflow scripts and the code for analyses and statistics are openly available on GitHub (https://github.com/MICA-MNI/2025_in-vivo_tauPET-mk6240_TLE).

## Acknowledgements

We thank the McConnell Brain Imaging Centre (BIC) PET and MRI units of the MNI, and the study participants for supporting our research. RRC received support from the Fonds de la Recherche du Québec – Santé (FRQ-S), the Montreal Neurological Institute Jeanne Timmins Costello Fellowship, and the Healthy Brains, Healthy Lives – Entrepreneur Postdoc Fellowship. JR received support from the Canadian Open Neuroscience Platform (CONP) and Canadian Institutes of Health Research (CIHR). MK is supported by the Wellcome Trust (221934/Z/20/Z). LC acknowledges previous support from Brain Research UK (Award 14181). BCB acknowledges support from CIHR (FDN-154298, PJT-174995), SickKids Foundation (NI17-039), Natural Sciences and Engineering Research Council (NSERC; Discovery-1304413), Azrieli Center for Autism Research of the Montreal Neurological Institute (ACAR), BrainCanada, FRQ-S, the Helmholtz International BigBrain Analytics and Learning Laboratory (Hiball), the Canada Research Chairs Program (CRC), and the Centre of Excellence in Epilepsy at the Neuro (CEEN).

## Supplementary Table 2

SUVR thresholds. Percentiles for SUVR values based on the mean cortical uptake of all vertices by group. Values represent standardized uptake value ratios (SUVR). We used the 97.5th percentile threshold based on healthy controls in the current manuscript.

## Glossary

TLE: temporal lobe epilepsy
PET: positron emission tomography
MRI: magnetic resonance imaging
NI: neuroinflammation

---

**Supplementary Figure 1.**
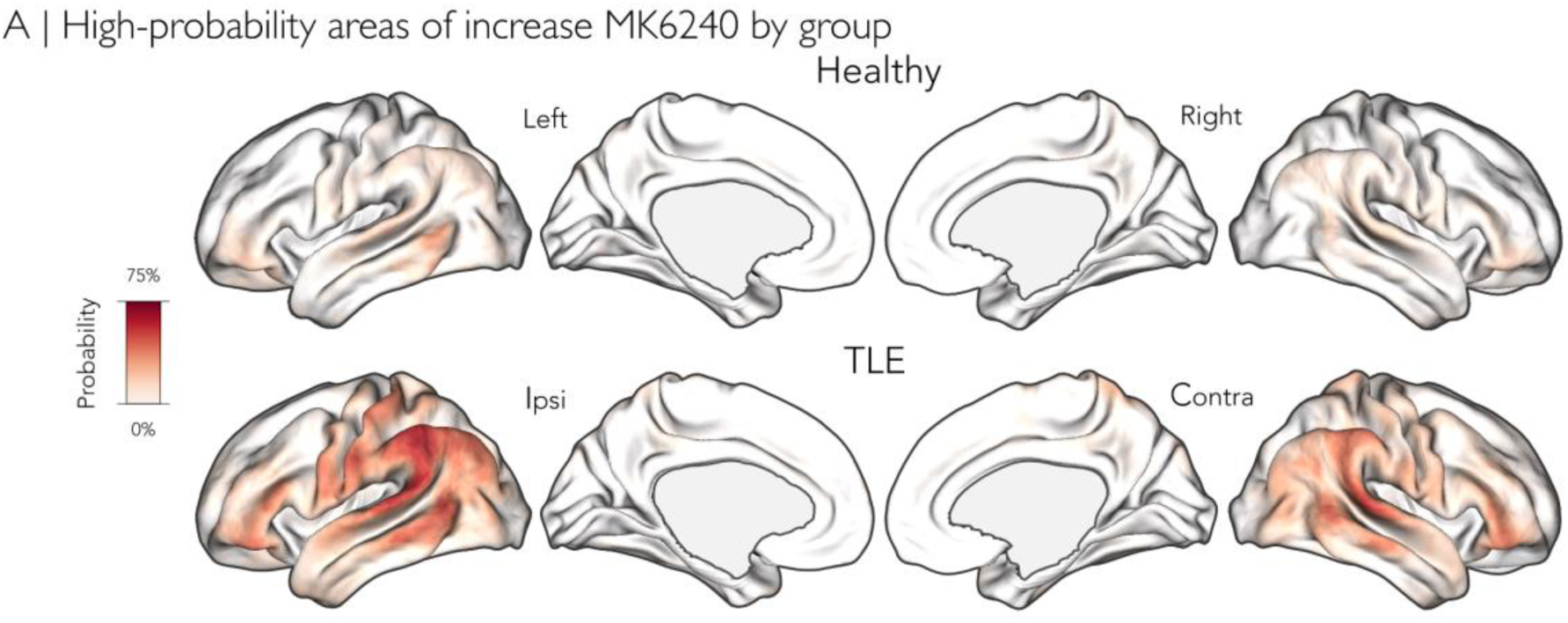
Cortical map of abnormal MK-6240 SUVR probability. Regions with SUVR values greater than 1.5 (percentile 97.5th based on controls) were identified and visualized for each group. Dark red represents areas with higher prevalence of abnormal SUVR uptake, light red indicates lower prevalence. The maximum MK-6240 SUVR probability is in 75% of the TLE population.

**Supplementary Figure 2.**
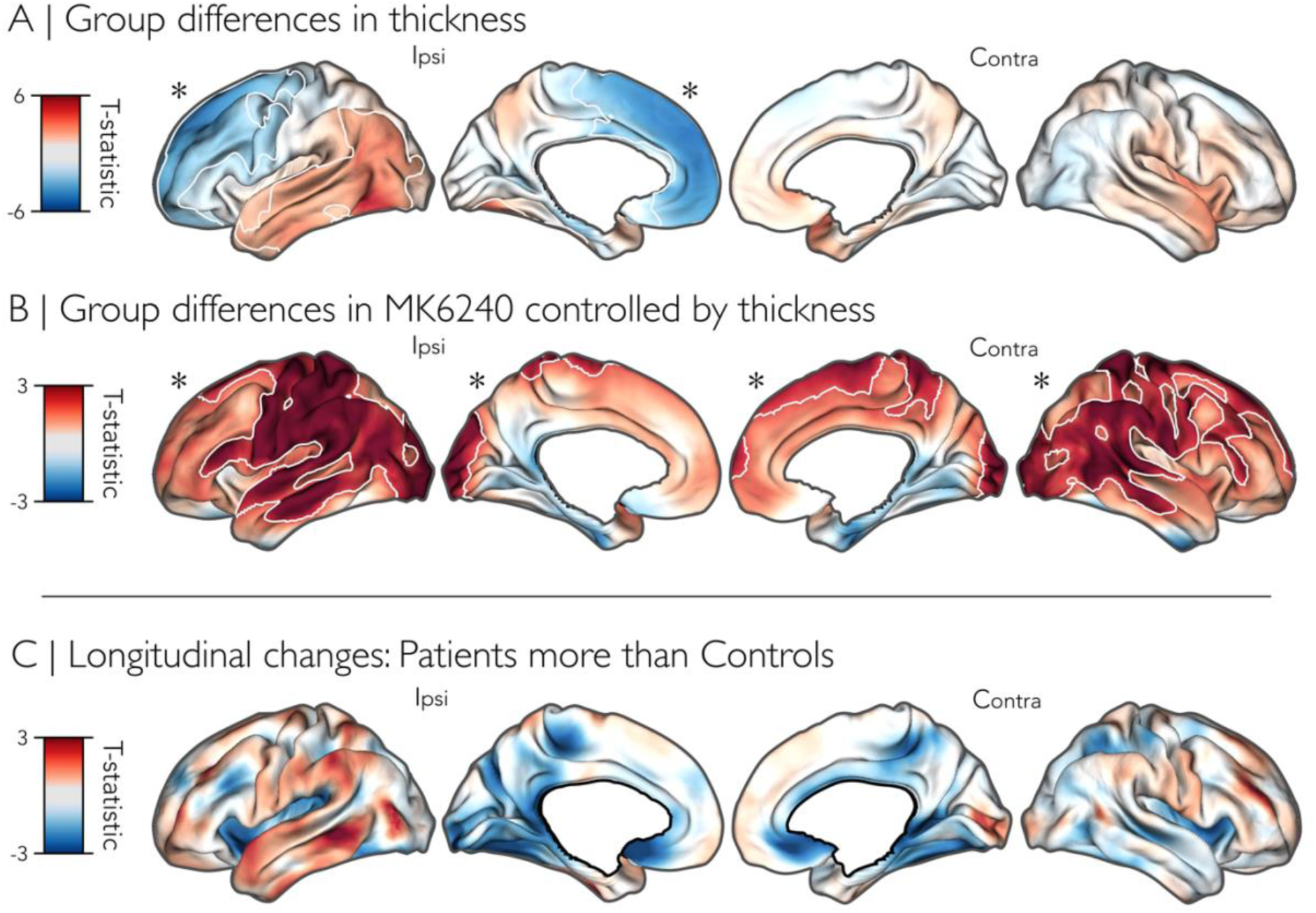
**A)** Impact of cortical thickness on between-group differences in MK-6240 uptake. Group difference in cortical thickness. The comparison was evaluated using a mixed effects model, controlling for age and sex, and corrected for multiple comparisons with random field theory (p<0.025). **B)** MK-6240 uptake comparing TLE and healthy controls after controlling for cortical thickness. **C)** Surface-based longitudinal changes in MK-6240 uptake comparing TLE and healthy controls. No cortical regions showed statistically significant longitudinal changes or group differences in the longitudinal slopes. Mixed-effects models were adjusted for age and sex and corrected for multiple comparisons using random field theory (p<0.025). Significant regions, when present, are outlined in white.

**Supplementary Figure 3.**
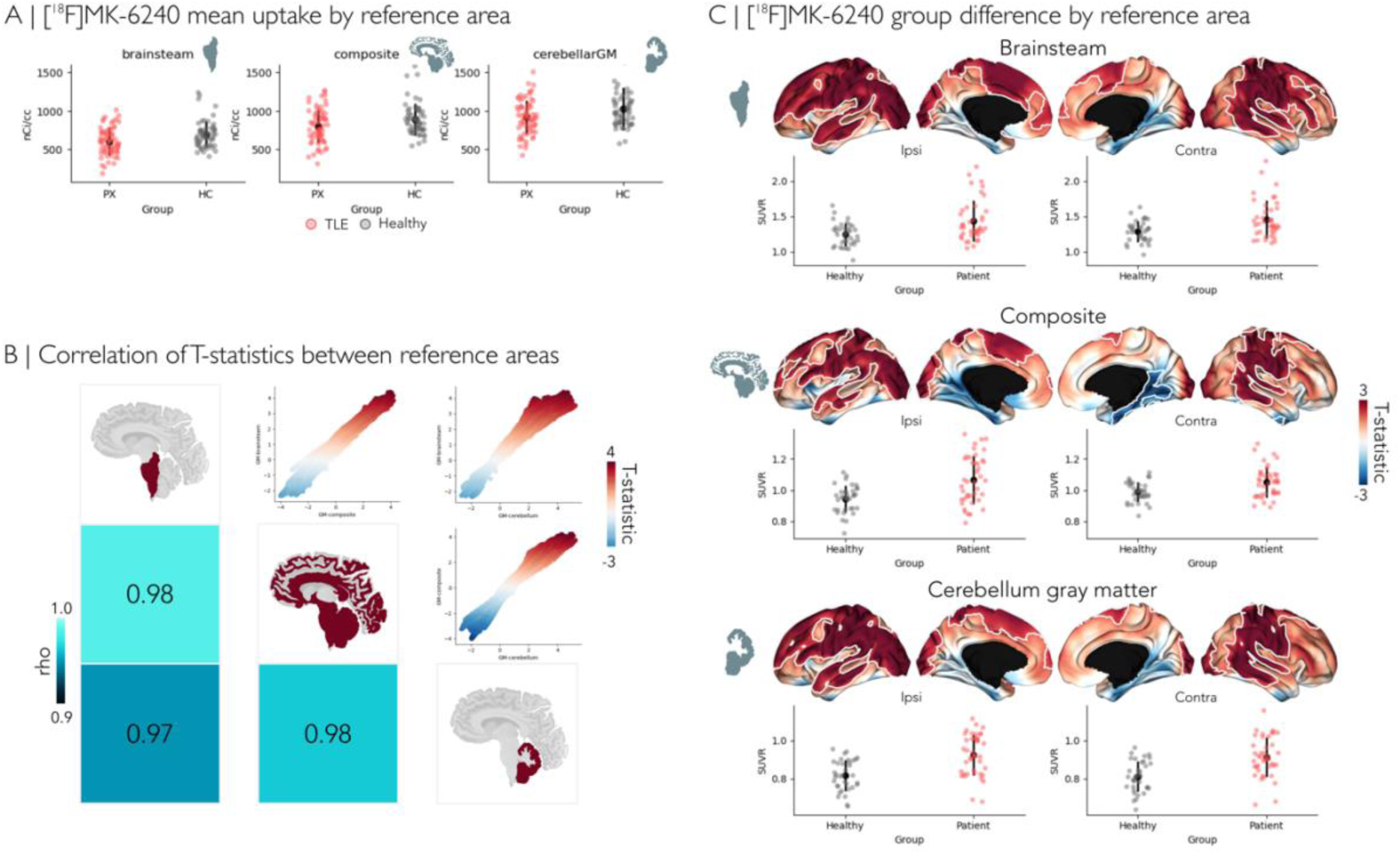
Consistency of MK-6240 quantification across reference regions. **A)** Scatterplot of the raw mean MK-6240 values in TLE patients and controls across different reference regions. **B)** Correlation of t-statistics from between-group difference maps across SUVR reference regions (brainstem, cerebellar gray matter, and a composite region). The top-right scatterplots show vertex-wise t-statistic comparisons for each pair of reference choices, along with Pearson correlation coefficients (r), demonstrating high consistency across methods (r ≥ 0.96). Blue-shaded brain maps illustrate the corresponding reference regions. The bottom-left matrix displays Spearman’s rho values between comparisons. **C)** Group comparisons between patients and controls for each reference region. The same mixed-effects model used in the main analyses was applied here, adjusting for age and sex, and corrected for multiple comparisons using random field theory (pRFT < 0.025). Significant cortical regions are outlined in white. Below the surfaces the scatter plots display mean SUVR values for significant regions in each hemisphere by group. Collectively, these results indicate strong agreement in regional t-statistics across the reference region choices used for MK-6240 SUVR computation.

**Supplementary Figure 4.**
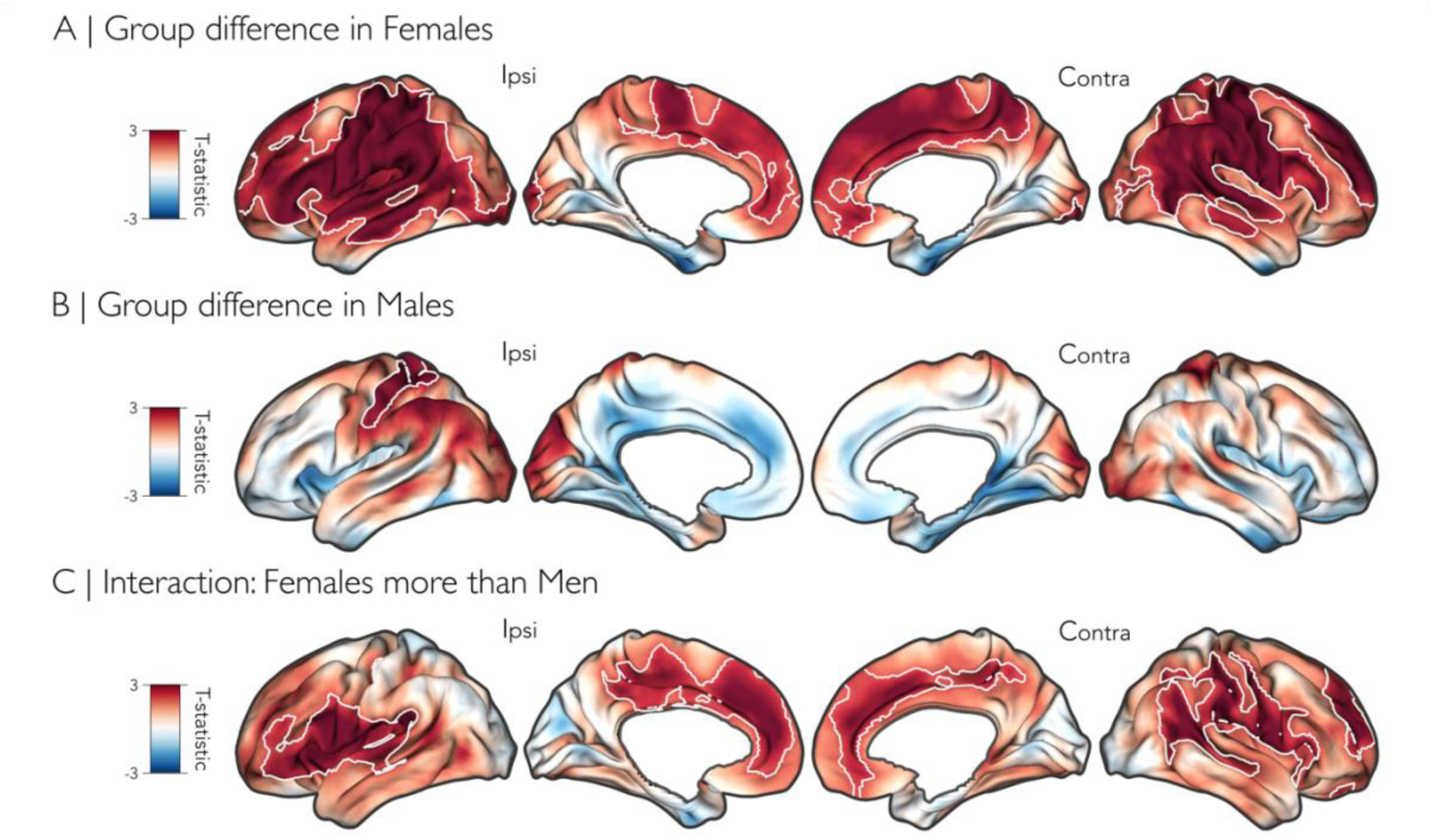
Sex differences in MK-6240 uptake. The comparison was evaluated using a mixed effects model, controlling for age, and corrected for multiple comparisons with random field theory (p<0.025). ***A)*** Group comparisons between only female patients and female controls. **B)** Group comparisons between only male patients and male controls**. C)** Group comparisons between patients and controls with an interaction of group by sex, where higher T-values represent where females have higher uptake than males.

**Supplementary Figure 5.**
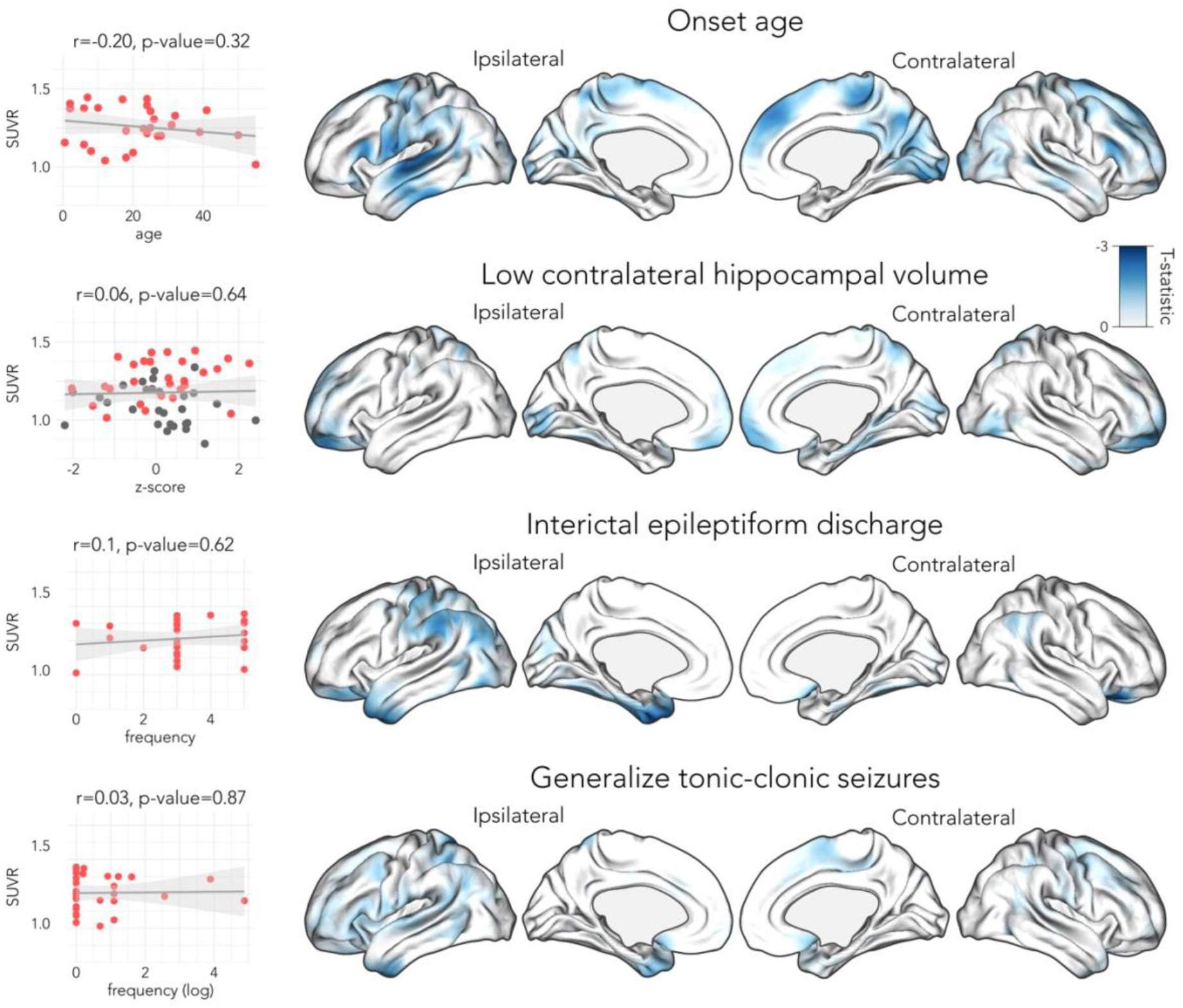
Tau MK-6240 SUVR and onset age, hippocampal volume, IEDs, GTCS and spatial task. Scatterplots display the relationship of mean significant MK-6240 SUVR with measures not explored in the main manuscript. For the spatial task only 12 out of 28 TLE patients had available data. Brain maps illustrate the corresponding regional effects, dark blue indicating areas where lower values of the corresponding measures are associated with higher MK-6240 SUVR. Mixed-effects models were used to assess these association, with correction for multiple comparisons with random field theory (p<0.025, two tails). No association reached statistical significance.

**Supplementary Figure 6.**
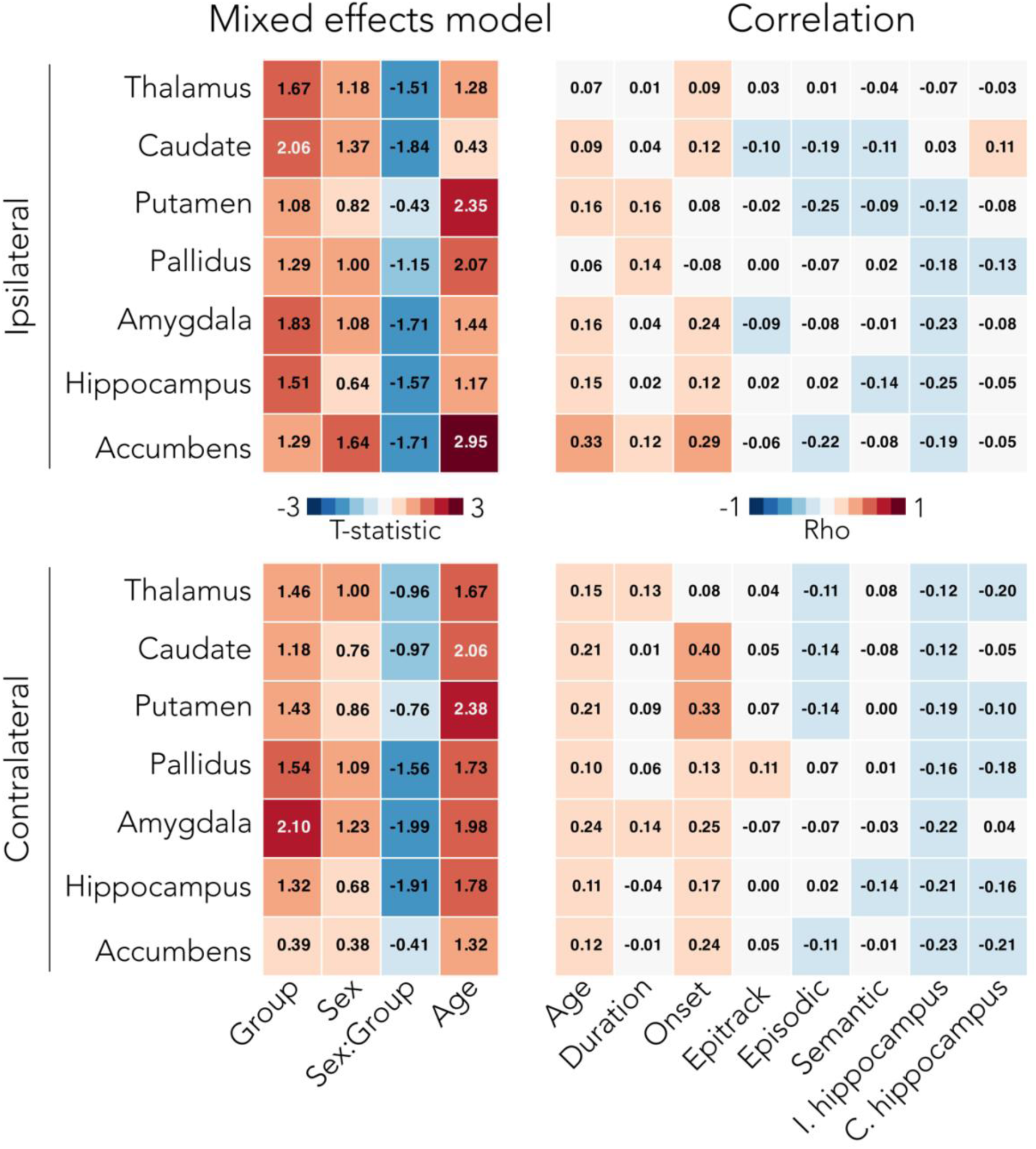
Group differences and correlations of subcortical and hippocampal MK-6240 uptake. The left panel shows the t-statistics from mixed-effects models examining group differences in each subcortical mean significant MK-6240 SUVR, including an interaction of group by sex and covariates: age, disease duration, and ipsilateral hippocampal volume (I.hippocampus). The right panel shows Spearman correlations between subcortical mean significant MK-6240 SUVRs and the same clinical and behavioral variables evaluated in the cortical MK-6240 analysis. P-values were corrected for multiple comparisons using the Bonferroni-Holm method; no results reached significance.

**Supplementary Figure 7.**
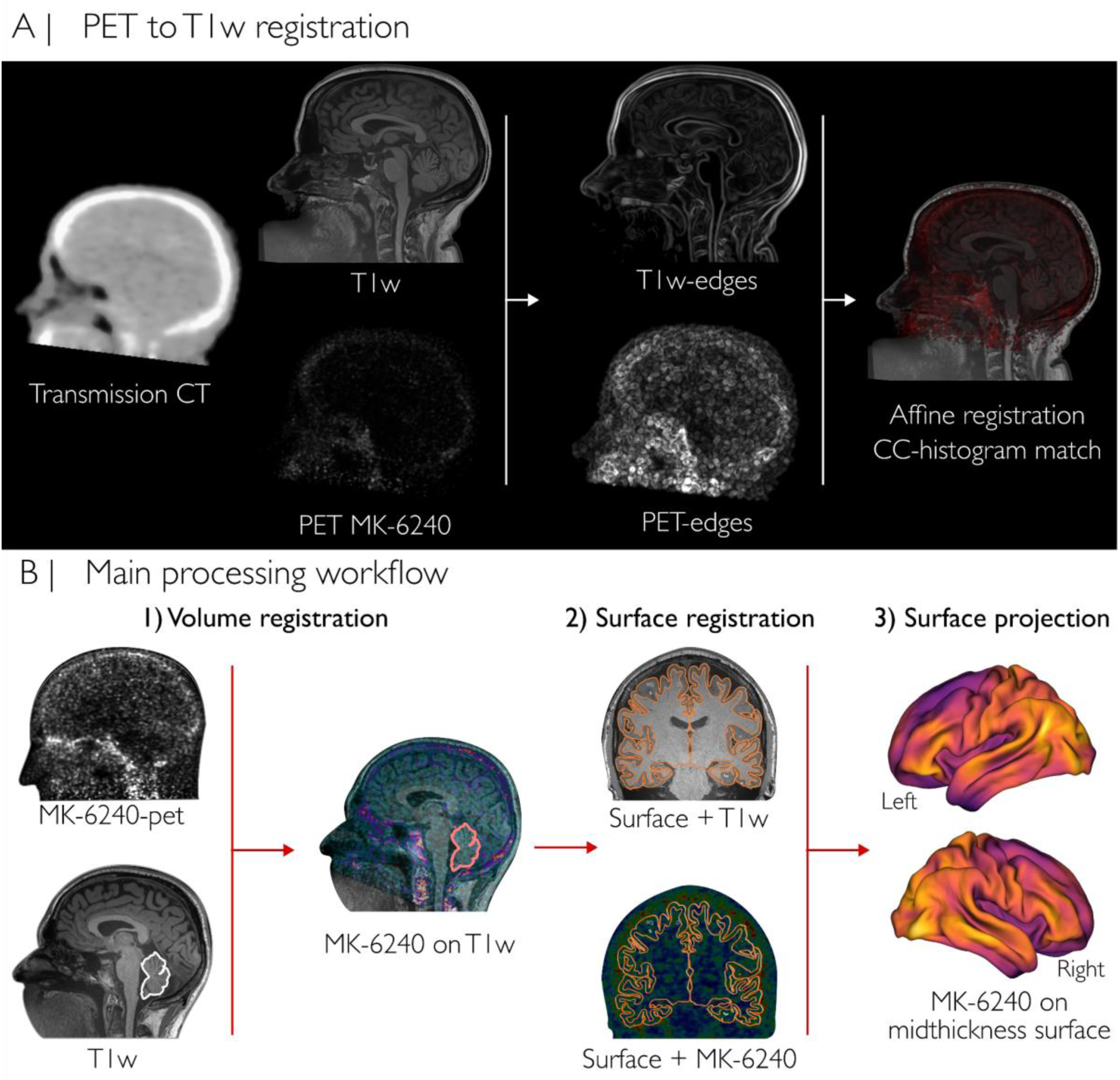
PET image processing workflow. **A)** Edge-based registration between PET MK-6240 and T1-weighted (T1w) MRI. Edge detection was applied to both PET and T1w images, and the resulting edge maps were affine-registered using a cross-correlation cost function with histogram matching. **B)** Workflow of co-registration between the structural T1w MRI and the MK-6240 PET image. Normalization by the cerebellar mean SUVR and mapping to the mid-thickness surface of each subject. The right image shows an example of one subject MK-6240 map projected on a standardized surface, fsLR-32k.

**Supplementary Figure 8.**
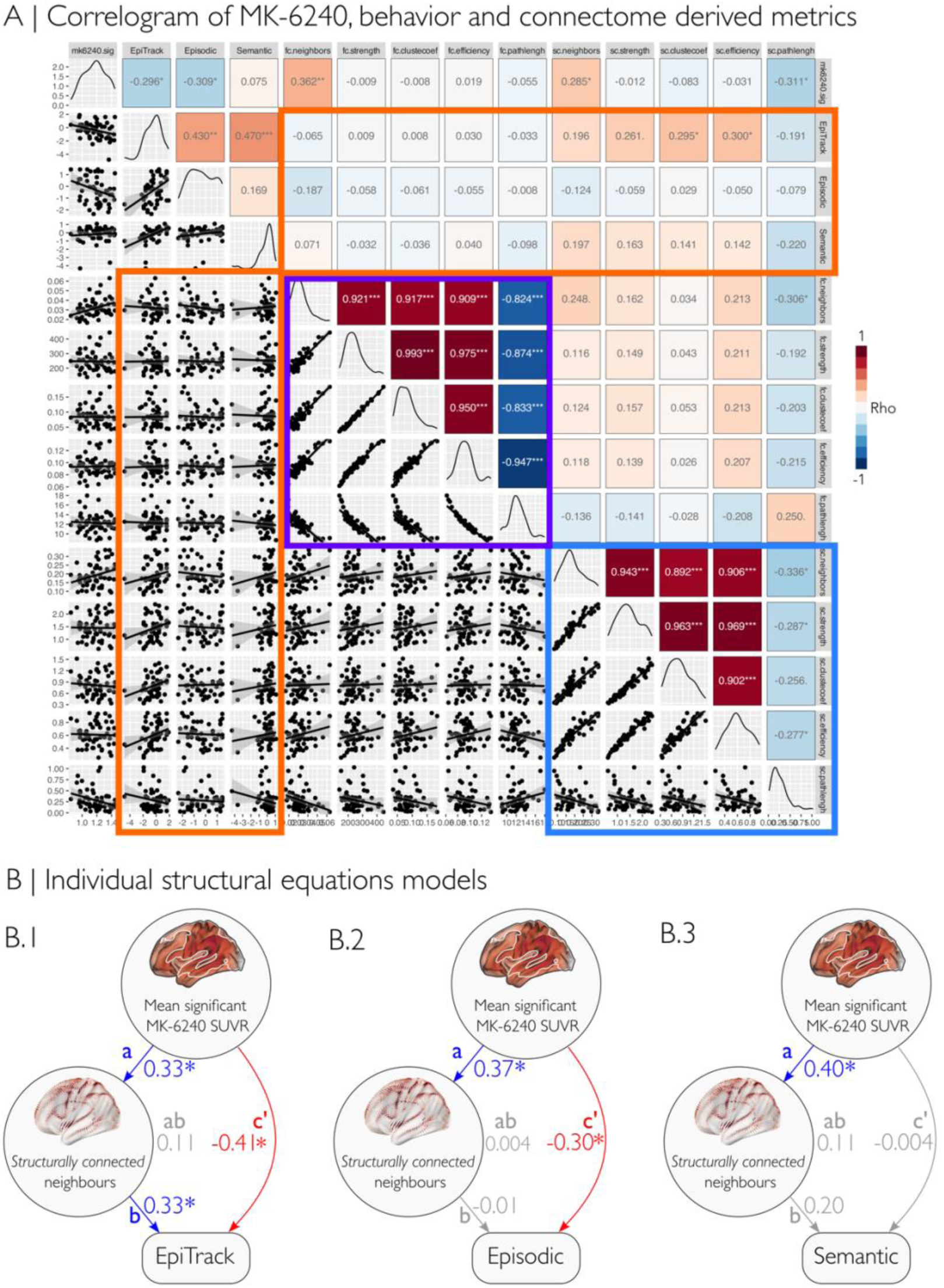
Impact of the connectome derived metric on the SUVR uptake and behavior. **A)** Correlogram of the connectome derived metrics, behavioral scores and mean MK-6240 from significant clusters. Orange square represents the correlations between behavioral scores and network derive metrics from structural and functional connectomes (neighborhood-weighted connectivity, strength, cluster coefficient, efficiency and characteristic path length). The purple rectangle in the center represents the correlations between network derived metrics from functional connectomes. The blue rectangle in the bottom-right represents the correlations between network derived metrics from structural connectomes. ***B)*** Individual Structural equation models showing relationships between mean significant MK-6240, structural network measures (SC weighted neighbors and SC efficiency), and behavioral outcomes. Values represent standardized path coefficients. Significant paths (p < 0.05) are shown in blue for positive effects and red for negative effects, the rest is gray. Indirect effects via semantic network measures were tested but were not significant.

**Supplementary Figure 9.**
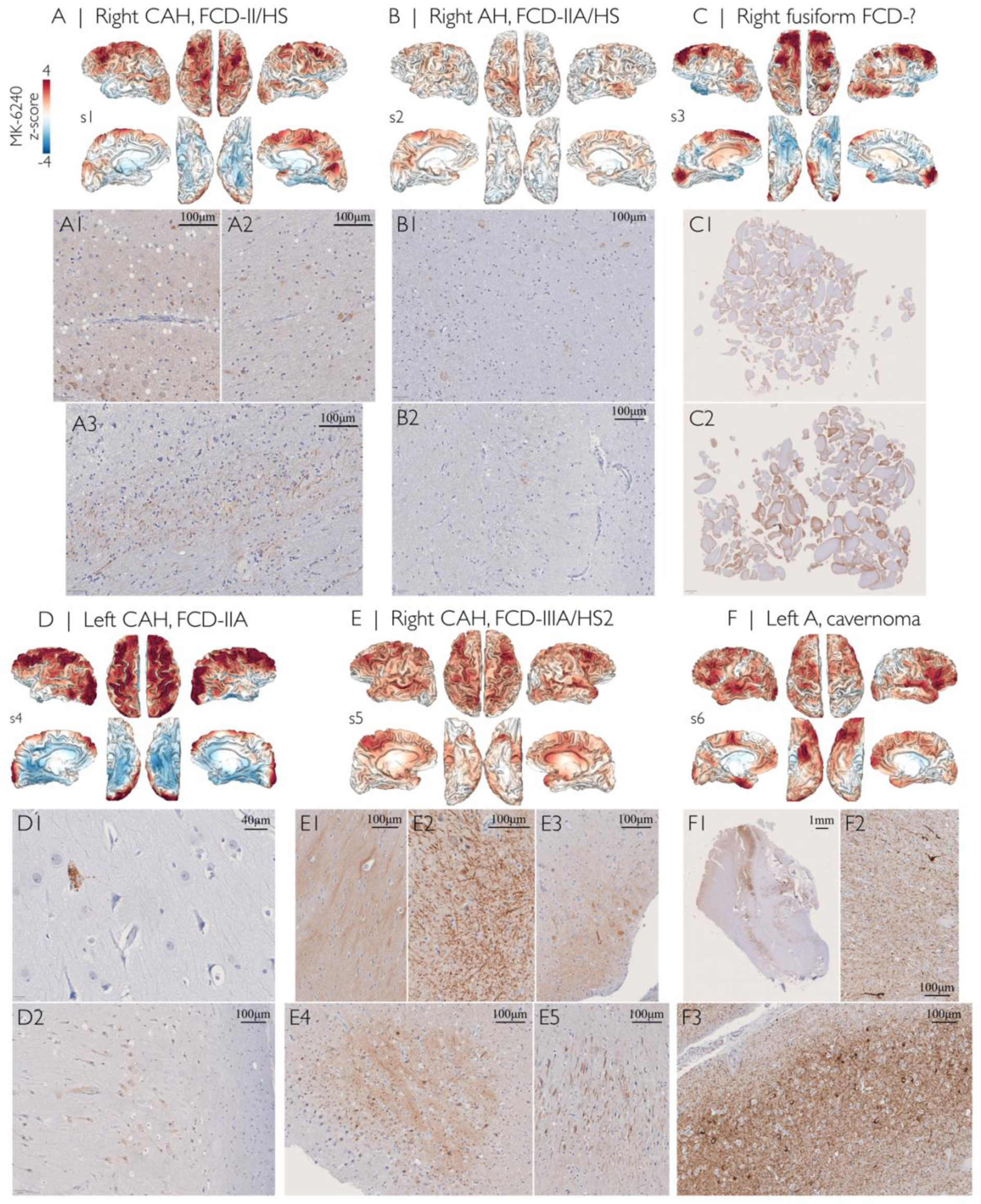
Cortical MK-6240 PET and AT8 immunohistochemistry in post-surgical tissue. Six cases are presented. For each case, cortical MK-6240 SUVR z-scores relative to controls are displayed at the top of each panel (range: −4 to 4). Cortical surface projections are shown in the following order (*left to right*): left lateral, left superior, right superior, and right lateral views (*top row*), followed by left medial, left inferior, right inferior, and right medial views (*bottom row*). Representative post-surgical tissue sections stained with AT8 immunohistochemistry are shown beneath each cortical surface. Panels correspond to individual cases labeled A–F, with multiple sections per case. (A1–A3, B1–B2, C1–C2, D1–D2, E1–E5, F1–F3).

## Notes

https://github.com/MICA-MNI/2025_in-vivo_tauPET-mk6240_TLE

https://osf.io/ct3gw

https://codeocean.com/capsule/4623457

### Competing Interest Statement

The authors have declared no competing interest.

### Summary of Updates

This version of the manuscript has been revised to include a subgroup analysis comparing patients with the highest SUVR uptake to the remaining cohort using cross-sectional data only. No substantial differences were observed in lateralization, hippocampal volumes, cognitive performance, seizure frequency, interictal epileptiform discharges, or hippocampal sclerosis. However, patients with higher SUVR uptake showed earlier age at onset and lower path length derived from structural connectomes.

## References

1. Pitkänen, A. & Sutula, T. P. Is epilepsy a progressive disorder? Prospects for new therapeutic approaches in temporal-lobe epilepsy. Lancet Neurol. 1, 173–181 (2002).

2. Galovic, M. et al. Progressive Cortical Thinning in Patients With Focal Epilepsy. JAMA Neurol 76, 1230–1239 (2019).

3. Tai, X. Y., et al. Review: Neurodegenerative processes in temporal lobe epilepsy with hippocampal sclerosis: Clinical, pathological and neuroimaging evidence. Neuropathol. Appl. Neurobiol. 44, 70–90 (2018).

4. Caciagli, L. et al. A meta-analysis on progressive atrophy in intractable temporal lobe epilepsy: Time is brain? Neurology 89, 506–516 (2017).

5. Boyle, P. A., Wilson, R. S., Aggarwal, N. T., Tang, Y. & Bennett, D. A. Mild cognitive impairment. Neurology 67, 441–445 (2006).

6. Noebels, J. A perfect storm: Converging paths of epilepsy and Alzheimer’s dementia intersect in the hippocampal formation. Epilepsia 52 **Suppl 1**, 39–46 (2011).

7. Lam, A. D. et al. Association of Seizure Foci and Location of Tau and Amyloid Deposition and Brain Atrophy in Patients With Alzheimer Disease and Seizures. Neurology 103, e209920 (2024).

8. Irizarry, M. C. et al. Incidence of new-onset seizures in mild to moderate Alzheimer disease. Arch. Neurol. 69, 368–372 (2012).

9. Amatniek, J. C. et al. Incidence and predictors of seizures in patients with Alzheimer’s disease. Epilepsia 47, 867–872 (2006).

10. Tai, X. Y. et al. Hyperphosphorylated tau in patients with refractory epilepsy correlates with cognitive decline: a study of temporal lobe resections. Brain 139, 2441–2455 (2016).

11. Salemme, S. et al. The prognosis of mild cognitive impairment: A systematic review and meta-analysis. Alzheimers Dement (Amst) 17, e70074 (2025).

12. Canet, G. et al. Seizure activity triggers tau hyperphosphorylation and amyloidogenic pathways. Epilepsia 63, 919–935 (2022).

13. Alves, M., Kenny, A., de Leo, G., Beamer, E. H. & Engel, T. Tau Phosphorylation in a Mouse Model of Temporal Lobe Epilepsy. Front. Aging Neurosci. 11, 490080 (2019).

14. Thom, M. & Koepp, M. Tau protein in drug-resistant epilepsy and cognitive decline. in Inflammation and Epilepsy: New Vistas 149–184 (Springer International Publishing, Cham, 2021).

15. Malarte, M.-L., Nordberg, A. & Lemoine, L. Characterization of MK6240, a tau PET tracer, in autopsy brain tissue from Alzheimer’s disease cases. Eur. J. Nucl. Med. Mol. Imaging 48, 1093–1102 (2021).

16. Aguero, C. et al. Autoradiography validation of novel tau PET tracer [F-18]-MK-6240 on human postmortem brain tissue. Acta Neuropathologica Communications 7, 1–15 (2019).

17. Hopewell, R. et al. A simplified radiosynthesis of [F]MK-6240 for tau PET imaging. J Labelled Comp Radiopharm 62, 109–114 (2019).

18. Knowlton, R. C. et al. In vivo hippocampal glucose metabolism in mesial temporal lobe epilepsy. Neurology 57, 1184–1190 (2001).

19. Juhász, C. et al. Glucose and [11C]flumazenil positron emission tomography abnormalities of thalamic nuclei in temporal lobe epilepsy. Neurology 53, 2037–2045 (1999).

20. Seeley, W. W., Crawford, R. K., Zhou, J., Miller, B. L. & Greicius, M. D. Neurodegenerative diseases target large-scale human brain networks. Neuron 62, 42–52 (2009).

21. de Bruin, H. et al. Connectivity as a universal predictor of tau progression in atypical Alzheimer’s disease. Brain 148, 3893–3912 (2025).

22. Ottoy, J. et al. Tau follows principal axes of functional and structural brain organization in Alzheimer’s disease. Nat Commun 15, 5031 (2024).

23. Kerestes, R. et al. Patterns of subregional cerebellar atrophy across epilepsy syndromes: An ENIGMA-Epilepsy study. Epilepsia 65, 1072–1091 (2024).

24. Whelan, C. D., Haaker, J. G. & Sisodiya, S. M. Structural Brain Abnormalities in the Common Epilepsies Assessed in a Worldwide ENIGMA Study. (2018).

25. Lutz, M. T. & Helmstaedter, C. EpiTrack: tracking cognitive side effects of medication on attention and executive functions in patients with epilepsy. Epilepsy Behav 7, 708–714 (2005).

26. Cabalo, D. G. et al. Differential reorganization of episodic and semantic memory systems in epilepsy-related mesiotemporal pathology. Brain 147, 3918–3932 (2024).

27. Tavakol, S. et al. Differential relational memory impairment in temporal lobe epilepsy. Epilepsy Behav 155, 109722 (2024).

28. Betthauser, T. J. et al. In Vivo Characterization and Quantification of Neurofibrillary Tau PET Radioligand 18F-MK-6240 in Humans from Alzheimer Disease Dementia to Young Controls. Journal of Nuclear Medicine 60, 93–99 (2019).

29. Pascoal, T. A. et al. 18F-MK-6240 PET for early and late detection of neurofibrillary tangles. Brain 143, 2818–2830 (2020).

30. Kreisl, W. C. et al. Patterns of tau pathology identified with 18F-MK-6240 PET imaging. Alzheimer’s & Dementia 18, 272–282 (2022).

31. Pascoal, T. A. et al. Longitudinal 18F-MK-6240 tau tangles accumulation follows Braak stages. Brain 144, 3517–3528 (2021).

32. Malpetti, M., La Joie, R. & Rabinovici, G. D. Tau Beats Amyloid in Predicting Brain Atrophy in Alzheimer Disease: Implications for Prognosis and Clinical Trials. Journal of Nuclear Medicine 63, 830–832 (2022).

33. Ossenkoppele, R. et al. Accuracy of Tau Positron Emission Tomography as a Prognostic Marker in Preclinical and Prodromal Alzheimer Disease: A Head-to-Head Comparison Against Amyloid Positron Emission Tomography and Magnetic Resonance Imaging. JAMA Neurol 78, 961–971 (2021).

34. La Joie, R. et al. Prospective longitudinal atrophy in Alzheimer’s disease correlates with the intensity and topography of baseline tau-PET. Sci. Transl. Med. 12, (2020).

35. Kunach, P. et al. Cryo-EM structure of Alzheimer’s disease tau filaments with PET ligand MK-6240. Nat Commun 15, 8497 (2024).

36. Gourmaud, S. et al. Alzheimer-like amyloid and tau alterations associated with cognitive deficit in temporal lobe epilepsy. Brain 143, 191–209 (2020).

37. Smith, K. M. et al. Tau deposition in young adults with drug-resistant focal epilepsy. Epilepsia 60, 2398–2403 (2019).

38. Thom, M. et al. Neurofibrillary tangle pathology and Braak staging in chronic epilepsy in relation to traumatic brain injury and hippocampal sclerosis: a post-mortem study. Brain 134, 2969–2981 (2011).

39. Ballerini, A. et al. Late-onset temporal lobe epilepsy: insights from brain atrophy and Alzheimer’s disease biomarkers. Brain 148, 185–198 (2025).

40. Bernasconi, N. Is epilepsy a curable neurodegenerative disease? Brain 139, 2336–2337 (2016).

41. Snyder, H. M. et al. Sex biology contributions to vulnerability to Alzheimer’s disease: A think tank convened by the Women’s Alzheimer’s Research Initiative. Alzheimers Dement 12, 1186–1196 (2016).

42. Wisch, J. K. et al. Sex-related Differences in Tau Positron Emission Tomography (PET) and the Effects of Hormone Therapy (HT). Alzheimer Dis Assoc Disord 35, 164–168 (2021).

43. Ferretti, M. T., Dimech, A. S. & Chadha, A. S. Sex and Gender Differences in Alzheimer’s Disease. (Academic Press, 2021).

44. Zeighami, Y., Tremblay, C. & Dadar, M. Sex differences in vulnerability to tau pathology: Impact on cognitive decline. Alzheimers Dement 21, e70634 (2025).

45. Lopez-Lee, C., Torres, E. R. S., Carling, G. & Gan, L. Mechanisms of sex differences in Alzheimer’s disease. Neuron 112, 1208–1221 (2024).

46. Hophing, L., Kyriakopoulos, P. & Bui, E. Sex and gender differences in epilepsy. Int Rev Neurobiol 164, 235–276 (2022).

47. Mrzyglod, A. et al. Patterns of phosphorylated tau accumulation in a spectrum of acquired and developmental brain lesions associated with refractory epilepsy. Epilepsia (2025) doi:10.1111/epi.18418.

48. Altmann, A. et al. A systems-level analysis highlights microglial activation as a modifying factor in common epilepsies. Neuropathol Appl Neurobiol 48, e12758 (2022).

49. Dickstein, L. P. et al. Neuroinflammation in neocortical epilepsy measured by PET imaging of translocator protein. Epilepsia 60, 1248–1254 (2019).

50. Langworth-Green, C. et al. Chronic effects of inflammation on tauopathies. Lancet Neurol 22, 430–442 (2023).

51. Toscano, E. C. B. et al. Hyperphosphorylated Tau in Mesial Temporal Lobe Epilepsy: a Neuropathological and Cognitive Study. Mol Neurobiol 60, 2174–2185 (2023).

52. O’Brien, T. J. et al. The utility of a 3-dimensional, large-field-of-view, sodium iodide crystal--based PET scanner in the presurgical evaluation of partial epilepsy. J Nucl Med 42, 1158–1165 (2001).

53. Theodore, W. H. et al. [18F]fluorodeoxyglucose positron emission tomography in refractory complex partial seizures. Ann Neurol 14, 429–437 (1983).

54. Vinton, A. B. et al. The extent of resection of FDG-PET hypometabolism relates to outcome of temporal lobectomy. Brain 130, 548–560 (2007).

55. Griffith, H. R. et al. Preoperative FDG-PET temporal lobe hypometabolism and verbal memory after temporal lobectomy. Neurology 54, 1161–1165 (2000).

56. Courtney, M. R. et al. F-FDG-PET hypometabolism as a predictor of favourable outcome in epilepsy surgery: protocol for a systematic review and meta-analysis. BMJ Open 12, e065440 (2022).

57. Sharpe, C. et al. Longitudinal changes of focal cortical glucose hypometabolism in adults with chronic drug resistant temporal lobe epilepsy. Brain Imaging Behav 15, 2795–2803 (2021).

58. Benedek, K., Juhász, C., Chugani, D. C., Muzik, O. & Chugani, H. T. Longitudinal changes in cortical glucose hypometabolism in children with intractable epilepsy. J Child Neurol 21, 26–31 (2006).

59. Zhou, J., Gennatas, E. D., Kramer, J. H., Miller, B. L. & Seeley, W. W. Predicting regional neurodegeneration from the healthy brain functional connectome. Neuron 73, 1216–1227 (2012).

60. Crossley, N. A. et al. The hubs of the human connectome are generally implicated in the anatomy of brain disorders. Brain 137, 2382–2395 (2014).

61. Royer, J. et al. Epilepsy and brain network hubs. Epilepsia 63, 537–550 (2022).

62. Georgiadis, F. et al. Connectome architecture shapes large-scale cortical alterations in schizophrenia: a worldwide ENIGMA study. Mol Psychiatry 29, 1869–1881 (2024).

63. Larivière, S. et al. Network-based atrophy modeling in the common epilepsies: A worldwide ENIGMA study. Sci Adv 6, (2020).

64. Liu, L. et al. Trans-synaptic spread of tau pathology in vivo. PLoS One 7, e31302 (2012).

65. Braak, H., Alafuzoff, I., Arzberger, T., Kretzschmar, H. & Del Tredici, K. Staging of Alzheimer disease-associated neurofibrillary pathology using paraffin sections and immunocytochemistry. Acta Neuropathol 112, 389–404 (2006).

66. Harrison, T. M. et al. Longitudinal tau accumulation and atrophy in aging and alzheimer disease. Ann Neurol 85, 229–240 (2019).

67. Pan, N., Liu, S., Ge, X., Zheng, Y. & for Alzheimer’s Disease Neuroimaging Initiative. Association of hippocampal atrophy with tau pathology of temporal regions in preclinical Alzheimer’s disease. J Alzheimers Dis 13872877251314785 (2025).

68. Liang, Y. et al. Association of apolipoprotein E genotypes with epilepsy risk: A systematic review and meta-analysis. Epilepsy Behav 98, 27–35 (2019).

69. Chen, Y. et al. Seizure frequency, APOE ε4, and cognitive function in older people with epilepsy. Acta Epileptol 7, 30 (2025).

70. Liu, S., He, Z., Shi, W. & Li, J. The association between APOE gene polymorphisms and the risk, characteristics, and prognosis of epilepsy: A systematic review and meta-analysis. Epilepsy Behav 160, 110070 (2024).

71. de Souza, L. C. et al. CSF tau markers are correlated with hippocampal volume in Alzheimer’s disease. Neurobiol Aging 33, 1253–1257 (2012).

72. Aschenbrenner, A. J., Gordon, B. A., Benzinger, T. L. S., Morris, J. C. & Hassenstab, J. J. Influence of tau PET, amyloid PET, and hippocampal volume on cognition in Alzheimer disease. Neurology 91, e859–e866 (2018).

73. Harrison, T. M. et al. Tau deposition is associated with functional isolation of the hippocampus in aging. Nat Commun 10, 4900 (2019).

74. Pan, F.-F. et al. Associations of hippocampal volumes, brain hypometabolism, and plasma NfL with amyloid, tau, and cognitive decline. Alzheimers Dement 21, e70005 (2025).

75. Bernhardt, B. C. et al. Temporal lobe epilepsy: Hippocampal pathology modulates connectome topology and controllability. Neurology 92, e2209–e2220 (2019).

76. Bernhardt, B. C. et al. The spectrum of structural and functional imaging abnormalities in temporal lobe epilepsy. Ann Neurol 80, 142–153 (2016).

77. Larivière, S. et al. Brain Networks for Cortical Atrophy and Responsive Neurostimulation in Temporal Lobe Epilepsy. JAMA Neurol 81, 1199–1209 (2024).

78. Simons, J. S., Graham, K. S. & Hodges, J. R. Perceptual and semantic contributions to episodic memory: evidence from semantic dementia and Alzheimer’s disease. J. Mem. Lang. 47, 197–213 (2002).

79. Pascoal, T. A. et al. In vivo quantification of neurofibrillary tangles with [F]MK-6240. Alzheimers. Res. Ther. 10, 74 (2018).

80. Landau, S. M. et al. Measurement of longitudinal β-amyloid change with 18F-florbetapir PET and standardized uptake value ratios. J Nucl Med 56, 567–574 (2015).

81. Van Essen, D. C., Glasser, M. F., Dierker, D. L., Harwell, J. & Coalson, T. Parcellations and hemispheric asymmetries of human cerebral cortex analyzed on surface-based atlases. Cereb. Cortex 22, 2241–2262 (2012).

82. Royer, J. et al. An Open MRI Dataset For Multiscale Neuroscience. Sci Data 9, 569 (2022).

83. Cruces, R. R. et al. Micapipe: A pipeline for multimodal neuroimaging and connectome analysis. Neuroimage 263, 119612 (2022).

84. Zhang, Y., Brady, M. & Smith, S. Segmentation of brain MR images through a hidden Markov random field model and the expectation-maximization algorithm. IEEE Trans. Med. Imaging 20, 45–57 (2001).

85. Woolrich, M. W. et al. Bayesian analysis of neuroimaging data in FSL. Neuroimage 45, S173–86 (2009).

86. Fischl, B. FreeSurfer. Neuroimage 62, 774–781 (2012).

87. Tournier, J.-D. et al. MRtrix3: A fast, flexible and open software framework for medical image processing and visualisation. Neuroimage 202, 116137 (2019).

88. Billot, B. et al. SynthSeg: Segmentation of brain MRI scans of any contrast and resolution without retraining. Med Image Anal 86, 102789 (2023).

89. Larivière, S. et al. BrainStat: A toolbox for brain-wide statistics and multimodal feature associations. Neuroimage 266, 119807 (2023).

90. Shafiei, G. et al. Spatial Patterning of Tissue Volume Loss in Schizophrenia Reflects Brain Network Architecture. Biol Psychiatry 87, 727–735 (2020).

91. Vos de Wael, R., et al. BrainSpace: a toolbox for the analysis of macroscale gradients in neuroimaging and connectomics datasets. Commun Biol 3, 103 (2020).

92. Alexander-Bloch, A. F. et al. On testing for spatial correspondence between maps of human brain structure and function. Neuroimage 178, 540–551 (2018).

93. Rubinov, M. & Sporns, O. Complex network measures of brain connectivity: uses and interpretations. Neuroimage 52, 1059–1069 (2010).

94. Barrat, A., Barthélemy, M., Pastor-Satorras, R. & Vespignani, A. The architecture of complex weighted networks. Proc Natl Acad Sci U S A 101, 3747–3752 (2004).

95. Opsahl, T. & Panzarasa, P. Clustering in weighted networks. Soc. Networks 31, 155–163 (2009).

96. Wasserman, S. & Faust, K. Social Network Analysis: Methods and Applications. (Cambridge University Press, 1994).

97. Luppi, A. I. et al. Systematic evaluation of fMRI data-processing pipelines for consistent functional connectomics. Nat Commun 15, 4745 (2024).

98. Zuo, X.-N., Biswal, B. B. & Poldrack, R. A. Reliability and Reproducibility in Functional Connectomics. (Frontiers Media SA, 2019).

